# Multiplexed Imaging Mass Cytometry of Chemokine Milieus in Metastatic Melanoma Characterizes Features of Response to Immunotherapy

**DOI:** 10.1101/2021.07.29.454093

**Authors:** Tobias Hoch, Daniel Schulz, Nils Eling, Julia Martínez Gómez, Mitchell P. Levesque, Bernd Bodenmiller

**Affiliations:** University of Zurich, Department of Quantitative Biomedicine, Zurich, 8057, Switzerland; ETH Zurich, Institute for Molecular Health Sciences, Zurich, 8093 Switzerland; Particles-Biology Interactions Laboratory, Empa, Swiss Federal Laboratories for Materials Science and Technology, St. Gallen, Switzerland; University Hospital Zurich, Department of Dermatology, Schlieren, 8952, Switzerland

## Abstract

Intratumoral immune cells are crucial for tumor control and anti-tumor responses during immunotherapy. Immune cell trafficking into tumors is mediated by chemokines, which are expressed and secreted upon various stimuli and interact with specific receptors. To broadly characterize chemokine expression and function in tumors, we have used multiplex mass cytometry-based imaging of protein markers and RNA transcripts to analyze the chemokine landscape and immune infiltration in metastatic melanoma samples. Tumors that lacked immune infiltration were devoid of most chemokines and exhibited particularly low levels of antigen presentation and inflammation. Infiltrated tumors were characterized by expression of multiple chemokines. *CXCL9* and *CXCL10* were often localized in patches associated with dysfunctional T cells expressing *CXCL13* which was strongly associated with B cell patches and follicles. TCF7^+^ naïve-like T cells, which predict response to immunotherapy, were enriched in the vicinity of B cell patches and follicles. Our data highlight the strength of RNA and protein co-detection which was critical to deconvolve specialized immune microenvironments in inflamed tumors based on chemokine expression. Our findings further suggest that the formation of tertiary lymphoid structures is accompanied by naïve and naive- like T cell recruitment, which ultimately boosts anti-tumor activity.

**One sentence summary:** Inflammatory chemokine milieus in metastatic melanoma are hotspots of T cell dysfunction and *CXCL13* expression, which likely guide the recruitment of B cells and the formation of B cell follicles responsible for anti-tumor immunity.

## INTRODUCTION

Immune checkpoint inhibition (ICI) has greatly improved the overall survival rates of patients with metastatic melanoma (*1, 2*); however, response rates vary, and factors that determine response to immunotherapy are largely unknown (*3*). Patients with inflamed (“immune hot”) tumors have better prognosis (*4*) and are more likely to benefit from ICI (*5*), whereas those with tumors with low levels of immune infiltration (“cold” or “deserted” tumors) are associated with lower ICI success (*6*). CD8^+^ T cells are critical mediators of anti-tumor immunity, and increased CD8^+^ T cell infiltration is associated with better prognosis in many tumor types (*4*). Important factors associated with CD8^+^ T cell infiltration are tumor antigen presentation and mutational burden (*7–9*). CD8^+^ T cell infiltration into tumors, followed by TCR stimulation via persistent antigen exposure, is often accompanied by CD8^+^ T cell dysfunction and exhaustion. This exhaustion is ideally reversed by ICI (*10, 11*). Currently, it is debated which CD8^+^ T cell phenotypes maintain anti-tumor immunity and which are most susceptible to reactivation via ICI (*12*). Two recently described T cell populations are the *CXCL13*-expressing dysfunctional CD8^+^ T cells, which are antigen-experienced and express high levels of exhaustion markers (*13, 14*), and the naive-like TCF7^+^ CD8^+^ T cells whose presence is associated with response to ICI in melanoma and non-small cell lung cancer (*12, 15–17*). Although the presence of B cells and tertiary lymphoid structures (TLS) has been shown to predict response to ICI in melanoma (*8, 18*), our knowledge about T cell dysfunction and the interplay with B cells in tumors remains incomplete.

Chemokines are small, secreted proteins that mediate immune cell trafficking (*19–21*) and immune cell recruitment into tumors (*22–24*). In particular, the expression of the CXCR3 ligands CXCL9 and CXCL10 by macrophages is required for the recruitment of CD8^+^ T cells into inflamed tumor tissues (*25–27*). CD8^+^ T cells exposed to cognate antigens may also express CCL4 to recruit additional CD8^+^ T cells via CCR5 (*28*). The chemokines CCL19 and CXCL13 function in T cell homing to lymph nodes via CCR7 and in B cell follicle maturation (*21*). A broad characterization of chemokine expression within the context of *in situ* tumors on the protein level is technically challenging and currently lacking.

Here, we harnessed the power of imaging mass cytometry (IMC) to study the chemokine landscape and additional factors that govern T cell infiltration and dysfunction in metastatic melanoma, with single-cell and spatial resolution. For this analysis, we extended our previously published RNA-protein co-staining protocol (*29, 30*) to enable robust detection of a dozen chemokines, for which antibodies are lacking or incompatible with IMC, expressed by tumor, immune, and stromal cells. The result is the first comprehensive chemokine map of metastatic melanoma. We show that chemokine-expressing cells are spatially organized in patches, that T cell-infiltrated tumors are characterized by strong chemokine expression and varying levels of T cell dysfunction, and that infiltration in melanoma is largely dependent on antigen presentation by tumor cells. We discovered that T cells are the sole source of *CXCL13* when B cells, but no B cell follicles, are present, suggesting that T cells drive the recruitment of B cells and potentially the formation of B cell follicles. Further, there is a spatial association between TLS and naive-like TCF7^+^ T cells, a cell type predictive of response to immunotherapy. We propose a model in which TCF7^+^ T cells emerge near B cell follicles to sustain anti-tumor reactivity.

## RESULTS

### Multiplexed RNA and protein detection with IMC

To study chemokine-dependent immune infiltration into tumors and to functionally characterize cell phenotypes, we extended our previously published RNA and protein co-stain protocol for IMC (*29*) to detect 12 mRNAs encoding chemokines and 29 proteins (Table S1). We used the 12 RNA channels to study the expression of mRNAs encoding chemokines with diverse functions in T cell attraction (*CXCL9*, *CXCL10*, *CXCL12*, *CXCL13*, *CCL2*, *CCL4*, *CCL18*, *CCL19*, *CCL22*), B cell attraction (*CXCL13*), neutrophil attraction (*CXCL8*), and monocyte, macrophage, and dendritic cell attraction (*CCL2*, *CCL4*, *CXCL12*). Probes to these RNAs as well as antibodies to detect additional 28 proteins were included in the “RNA & protein” panel. The “protein” panel contained markers to identify and characterize tumor cells (Sox9, Sox10, MITF, Ki-67, S100A1, p75, β-Catenin, H3K27me3, pERK, pS6, PD-L1) and additional immune cell types and phenotypes (e.g., CD303, CD20, GrzB, PD-1, TCF7). Together, these panels enabled characterization and spatial analysis of tumor phenotypes in hot and cold tumors, of the tumor immune microenvironment, and of T cell phenotypes.

In experiments to validate mRNA detection, signal intensities across all 12 channels for detection of chemokine-expressing mRNAs were comparable to the signal intensity for a control *PPIB* gene (Fig. S1A), and expression levels of the 12 housekeeping genes measured with the IMC mRNA method highly correlated to those determined by bulk RNA sequencing (Pearson’s r = 0.78; Fig. S1B). We found no differences in signal intensities for the RNA probes with and without subsequent antibody staining (Fig. S1C), and an assessment of cross-hybridization between channels showed that there was minimal crosstalk (maximum of 2.1% between channels; Fig. S1D, E). Thus, we concluded that the 12-plex mRNA detection is compatible with multiplex protein IMC with negligible channel crosstalk at endogenous levels of expression.

### Heterogeneous immune infiltration in metastatic melanoma

We applied our multiplexed RNA and protein IMC panel to study immune infiltration in metastatic melanoma. The tissue microarray (TMA) we analyzed consisted of multiple cores from formalin-fixed paraffin-embedded (FFPE) tissue from 69 patients. The samples were from different metastatic sites, and cancers ranged in grade from Stage III to IV (Fig. 1A; Fig. S1F). Consecutive sections of the TMA were stained with the two panels, and images were acquired by IMC (Fig. 1A).

**Fig. 1.**
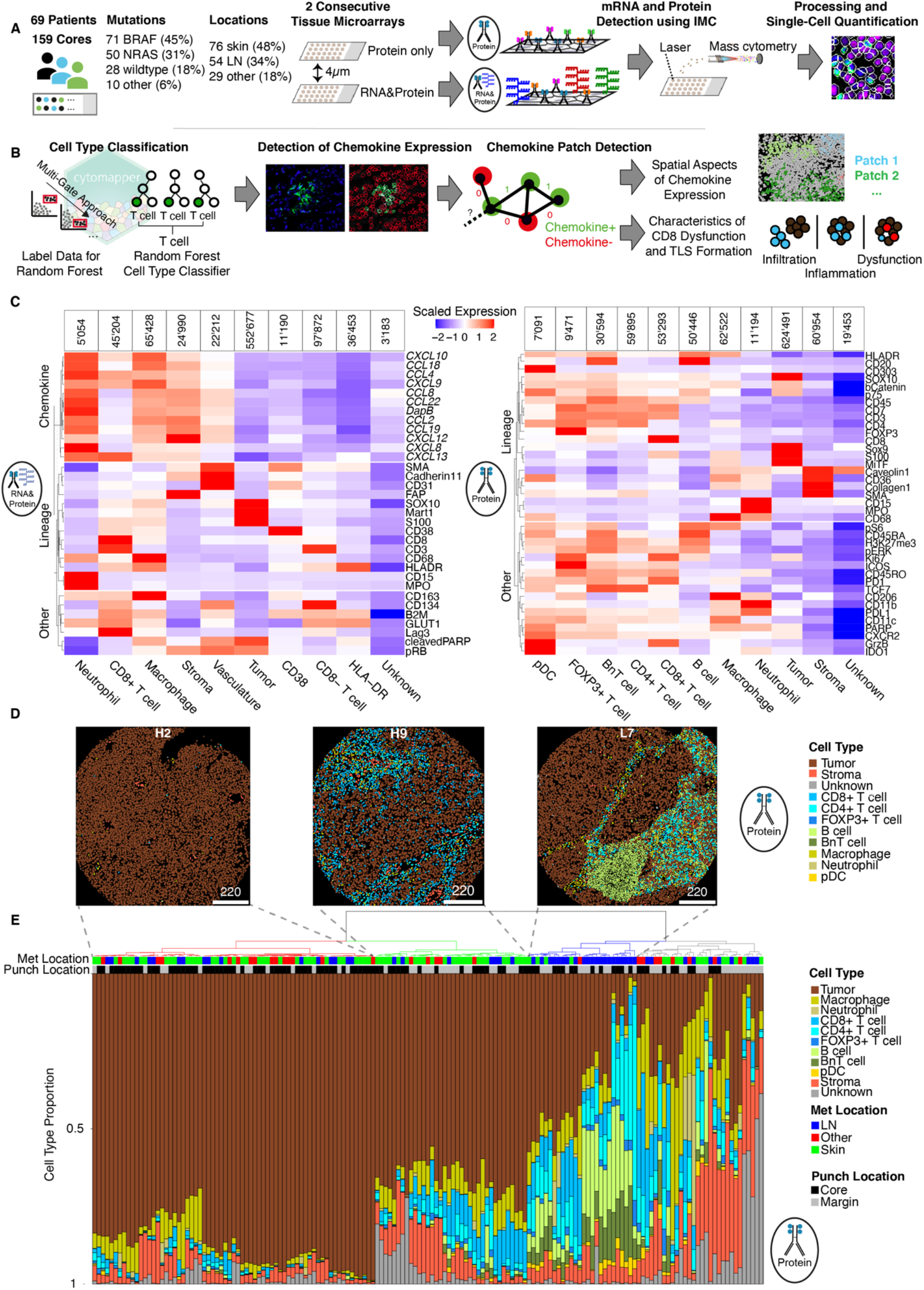
Profiling metastatic melanoma with RNA and protein co-detection using IMC. **(A)** Schematic of the IMC data acquisition of two consecutive slices of a TMA containing 159 biopsy cores from a total of 69 metastatic melanoma patients. Samples were stained with the RNA & protein panel and with the protein panel. Symbols for RNA & protein and protein panels are used to indicate the source of the data in subsequent figures. **(B)** Schematic of the computational workflow applied to the single-cell data. A random forest classifier was used to identify major cell types. A method for robust detection of cells that express chemokine mRNAs was used to assign chemokine-expressing cells. Cell phenotype and spatial aspects from both datasets were combined and used to characterize immune infiltration and T cell dysfunction in the melanoma samples. **(C)** Heat maps depicting scaled expression of markers in defined cell types of the RNA & protein dataset (left) and protein dataset (right). The number total number of cells of each cell type is indicated above each column. **(D)** Cell masks colored by cell type as identified in the protein panel for three different patient samples demonstrating heterogeneity across samples. **(E)** Stacked bar plot clustered by fractions of each cell type in single images. The metastasis locations and biopsy punch locations are shown as annotations on top of the bar plot. Images were hierarchically clustered using Ward’s method on Euclidean distances. The four major branches are indicated by color: red, very low frequencies of immune cells; green, moderate frequencies of immune cells; blue, high frequencies of lymphocytes enriched in lymph node samples; gray, high frequencies of immune and stromal cells and low frequencies of B cells.

We carried out single-cell segmentation and spill-over compensation to obtain single-cell data. We identified major cell types by supervised cell type labeling using *cytomapper* (*31*) followed by random forest classification. Data on 864,263 and 989,404 cells were obtained using the RNA & protein and protein staining panels, respectively (Fig. 1B). The most abundant cell populations were tumor cells (64.7%). There were similar frequencies of macrophages (6.5%), CD8^+^ T cells (5.5%), and CD4^+^ T cells (5.6%). We also observed a substantial fraction of B cells (4.1%) as well as densely packed cells that were indistinguishable from B or T cells (referred to here as BnT cells; 2.7%). Neutrophils (1.1%), T regulatory cells (1%), and plasmacytoid dendritic cells (0.7%) were the least abundant cell types (Fig. 1C). For all cell types that could be identified in both the RNA & protein dataset and the protein dataset we obtained a very high mean correlation of the cell-type frequencies (Pearson’s r = 0.94; Fig. S1G).

We then investigated whether our multiplex imaging captures the diverse landscapes of hot and cold metastatic melanomas. There were considerable differences in tumor and immune cell abundances across samples in the cohort when individual images were inspected (Fig. 1D). Clustering of the cell type frequencies from individual images revealed four major groups that reflect different levels of immune cell infiltration (Fig. 1E): images with low frequencies of immune cells, images with moderate frequencies of immune cells, images with high numbers of lymphocytes enriched in lymph node metastasis samples, and images dominated by immune and stromal cells with low frequencies of B cells.

### Chemokine expression in metastatic melanoma

To further investigate the basis of the observed heterogeneity of immune infiltration into tumors, we examined chemokine expression across the cohort. Chemokines showed distinct spatial expression patterns, were expressed by various cell types, and were globally detected via negative control normalization (Fig. 2A, Fig. S1H). Overall, tumor cells accounted for 22.4% of chemokine-expressing cells, followed by macrophages (21.9%), CD3^+^/CD8^-^ T cells (21%), and CD3^+^/CD8^+^ T cells (16.3%). However, only 2.4% of all tumor cells expressed chemokines, and only rarely did more than 10% of the tumor cells in a given image express chemokines (Fig. S2A). Of all cells in the RNA & protein dataset, 6.9% were chemokine expressing. Some of these cells expressed two chemokines (16.3% of all chemokine-expressing cells) or three or more chemokines (4.8% of chemokine-expressing cells).

**Fig. 2.**
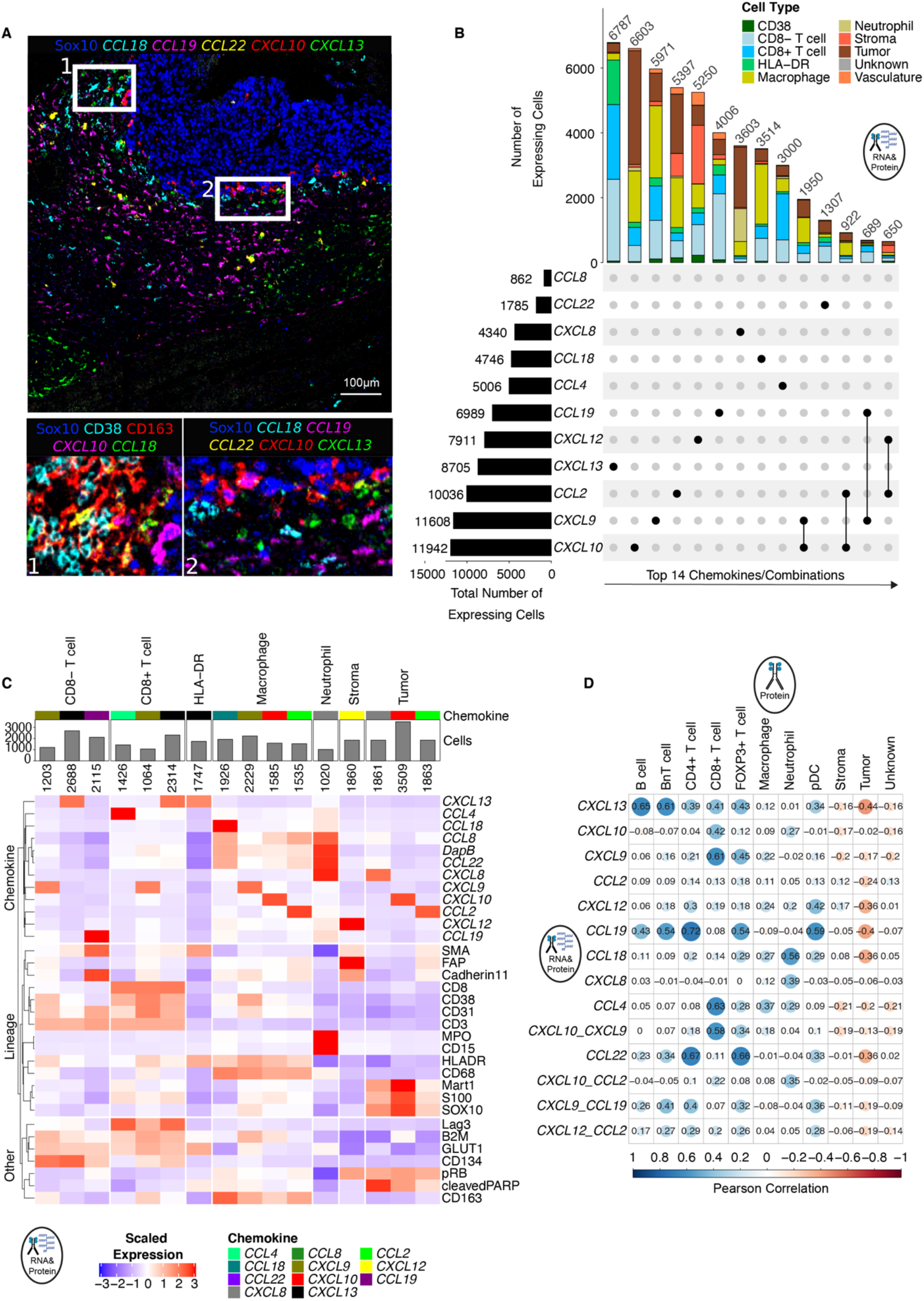
Chemokine expression landscape in metastatic melanoma. **(A)** Representative IMC image showing the expression of multiple chemokine mRNAs and tumor-marker Sox10. Box regions marked 1 and 2 are magnified below. Marker expression was false colored, and markers are indicated above each plot. A Gaussian blur (sigma = 0.65) was applied. Scale bar, 100 µm. **(B)** Upper stacked bar plot of the number of cells of indicated cell types that express the chemokine or chemokine combinations indicated by points below the stacked bar plot. The bar plot on the bottom left shows the total number of cells positive for the indicated chemokine. **(C)** Heat map of scaled expression of markers in chemokine-expressing cells with a minimal abundance of 1000 cells. Absolute cell numbers are shown on top of the heat map and with a bar plot. The colored boxes on top code for the chemokine which is expressed by a given cell type. **(D)** Pearson correlations between the frequencies of the most abundant chemokine combinations and frequencies of cell types from consecutive sections.

The most frequently expressed chemokine in singly-expressing cells was *CXCL13* and T cells accounted for a large fraction (71.1%) of cells that expressed *CXCL13* (Fig. 2B). *CXCL9* and *CXCL10* were the most frequently expressed chemokines, when cells that also expressed more than one chemokine were evaluated, followed by *CCL2* and *CXCL13* (Fig. 2B). Macrophages frequently expressed *CCL2*, *CCL18*, *CXCL9*, and *CXCL10*. *CXCL8* was mostly expressed by tumor cells and neutrophils. Stromal cells were a minority of chemokine-expressing cells (3.4%); stromal cells that did express a chemokine expressed *CXCL12* and weakly *CCL2*. *CCL8* was rarely and weakly expressed (862 cells total).

Chemokines are mostly expressed as an immediate response to external stimuli, such as inflammatory signals, and can therefore be used to derive phenotypic states. We examined mean marker expression in various cell types as a function of chemokine expression (Fig. 2C). CD8^+^ T cells were the strongest expressors of *CCL4*, and this subset of cells also highly expressed the exhaustion marker Lag-3 (Fig. 2C). Thus, these cells may represent a recently suggested antigen experienced “recruiter” state (*28*). Another subset of CD8^+^ T cells showed prominent *CXCL13* expression and also expressed Lag-3 (Fig. 2C). These cells are likely the recently reported “dysfunctional”, antigen-experienced CD8^+^ T cells (*13, 14, 32*). Of note, *CXCL9* and *CCL19* expression by T cells (CD8^+^ and CD8^-^) was most likely due to imperfect segmentation with either HLA-DR^+^ neighboring myeloid cells, which are the main cell type that express *CXCL9*, or *CCL19*-expressing fibroblastic reticular cells for which we do not have additional markers (Fig. S2B). Neutrophils were the strongest expressors of *CXCL8*, which has not been previously reported. Macrophages expressing *CCL18* expressed high levels of the M2 macrophage marker CD163. CD38 was detected at low levels on *CXCL9*-expressing macrophages, suggesting that these macrophages may have been activated by IFN-ɣ (*33*).

Other than the lymphoid tissue-specific chemokines *CXCL13* and *CCL19*, which were expressed at higher frequencies in lymph node metastasis samples than in subcutaneous skin metastasis samples (Fig. S2C), chemokine expression was similar in samples from subcutaneous skin and from lymph nodes. We also compared chemokine expression in samples from patients with *BRAF* and *NRAS* mutations. We observed elevated frequencies of chemokine-expressing cells, especially for inflammatory chemokines *CXCL9*, *CXCL10*, and *CCL2*, in samples with *BRAF* mutations compared to *NRAS* mutation and wild-type samples (Fig. S2D).

In summary, the detection of chemokine expression in tissues coupled with cell-type identification allowed us to quantify the chemokine expression landscape in metastatic melanoma. At the RNA level, chemokine expression was predominantly observed in immune cells. Furthermore, it was rare that cells expressed more than one chemokine.

### Chemokine expression-associated immune infiltration landscape

To identify potential chemokine-driven immune cell type recruitment we investigated the co-occurrence of cell types and chemokine-expressing cells, evaluating immune cell type frequencies from the protein dataset and the frequencies of chemokine-expressing cells in consecutive sections from the RNA & protein dataset (Fig. 2D). We observed moderate correlations between *CXCL13* and B cells and BnT cells (Pearson’s r = 0.65 and 0.61, respectively) in agreement with the importance of *CXCL13* for B cell recruitment. The frequencies of *CXCL9*- and *CCL4*-expressing cells correlated moderately with CD8^+^ T cells (Pearson’s r = 0.61 and 0.63, respectively). *CXCL9* was more highly correlated with the presence of CD8^+^ T cells than was *CXCL10* (Pearson’s r = 0.61 and 0.42, respectively), suggesting a dominant role for *CXCL9* in CD8^+^ T cell recruitment via CXCR3. The frequency of CD4^+^ T cells was not correlated with *CXCL9* or *CXCL10* but was moderately to highly correlated with *CCL22* and *CCL19* (Pearson’s r = 0.67 and 0.72, respectively), suggesting recruitment via CCR4 and CCR7, respectively. The frequency of regulatory T cells correlated most strongly with *CCL22* and moderately with *CCL19* (Pearson’s r = 0.66 and 0.54, respectively) in line with recruitment via CCR4 and CCR7. There was a low correlation between plasmacytoid dendritic cell frequencies and the presence of *CXCL12*-expressing cells (Pearson’s r = 0.42) and a moderate correlation with the presence of *CCL19*-expressing cells (Pearson’s r = 0.59), in agreement with plasmacytoid dendritic cell chemotaxis through CXCR4 and CCR7, respectively (*34, 35*). Although the CXCL8-CXCR1 axis is important for recruitment of neutrophils (*36*), the frequency of *CXCL8*-expressing cells was poorly correlated with the frequency of neutrophils in our dataset (Pearson’s r = 0.39). The chemotactic function of CCL2 toward multiple immune cell types has been reported (*37*); however, the frequency of *CCL2*-expressing cells in our dataset was not correlated with any other cell type frequency. The chemokine encoding mRNA that showed strongest correlation with the frequency of macrophages was *CCL4*, and this correlation was low (Pearson’s r = 0.37).

Only a subset of the chemokines showed co-expression in images, and these were mostly the inflammatory response chemokines *CXCL9*, *CXCL10*, *CCL4*, and *CXCL13* (Fig. S2E). The presence of *CCL19*-expressing cells was moderately correlated with the presence of *CCL22*-expressing cells (Pearson’s r = 0.68) in line with functions for T cell homing to B cell follicles (*38*). Finally, we did not find a negative correlation amongst any chemokine-expressing cells, indicating an overall positive role in attraction and co-regulation.

Samples from our cohort showed considerable heterogeneity with respect to overall immune cell infiltration (Fig. 1F). Since the abundance of CD8^+^ T cells is prognostic for long-term patient survival (*4*), we focused more deeply on the association of chemokine-expressing cells with CD8^+^ T cell infiltration. To reflect hot and cold properties of tumors we grouped images based on their T cell densities, resulting in four groups, and compared the fractions of all chemokine-expressing cells. The fractions of all chemokine-expressing cells, except for those that expressed *CXCL8*, increased with T cell density (Fig. S3A). The strongest increases were observed for cells expressing *CCL4*, *CXCL9*, *CXCL10*, *CXCL12*, and *CXCL13*, similar to what has been previously shown (*22, 23*). Notably, *CXCL9* was virtually absent from regions lacking CD8^+^ T cells underlining the importance of *CXCL9* in CD8^+^ T cell recruitment.

### Chemokine-expressing cells form specialized milieus

We next asked whether chemokines are expressed in a spatially coordinated manner and thus show patterns of local enrichment. We performed a local enrichment analysis for all chemokines and observed that *CXCL10* was prominently expressed in clusters of neighboring cells compared to a random cell localization (Fig. 3A, S3B), similar to our previous observations in breast cancer (*29*). Over a third (36%) of all images were locally enrichment for four *CXCL10*-expressing neighboring cells. *CXCL9*- and *CXCL13*-expressing cells also showed frequent local enrichment of varying numbers of cells. Conversely, cells expressing chemokines *CCL4* and *CXCL8* were rarely locally enriched (3% and 7.2% of images for local enrichment of four cells, respectively). These findings suggest that some chemokines are subject to a coordinated expression program.

**Fig. 3.**
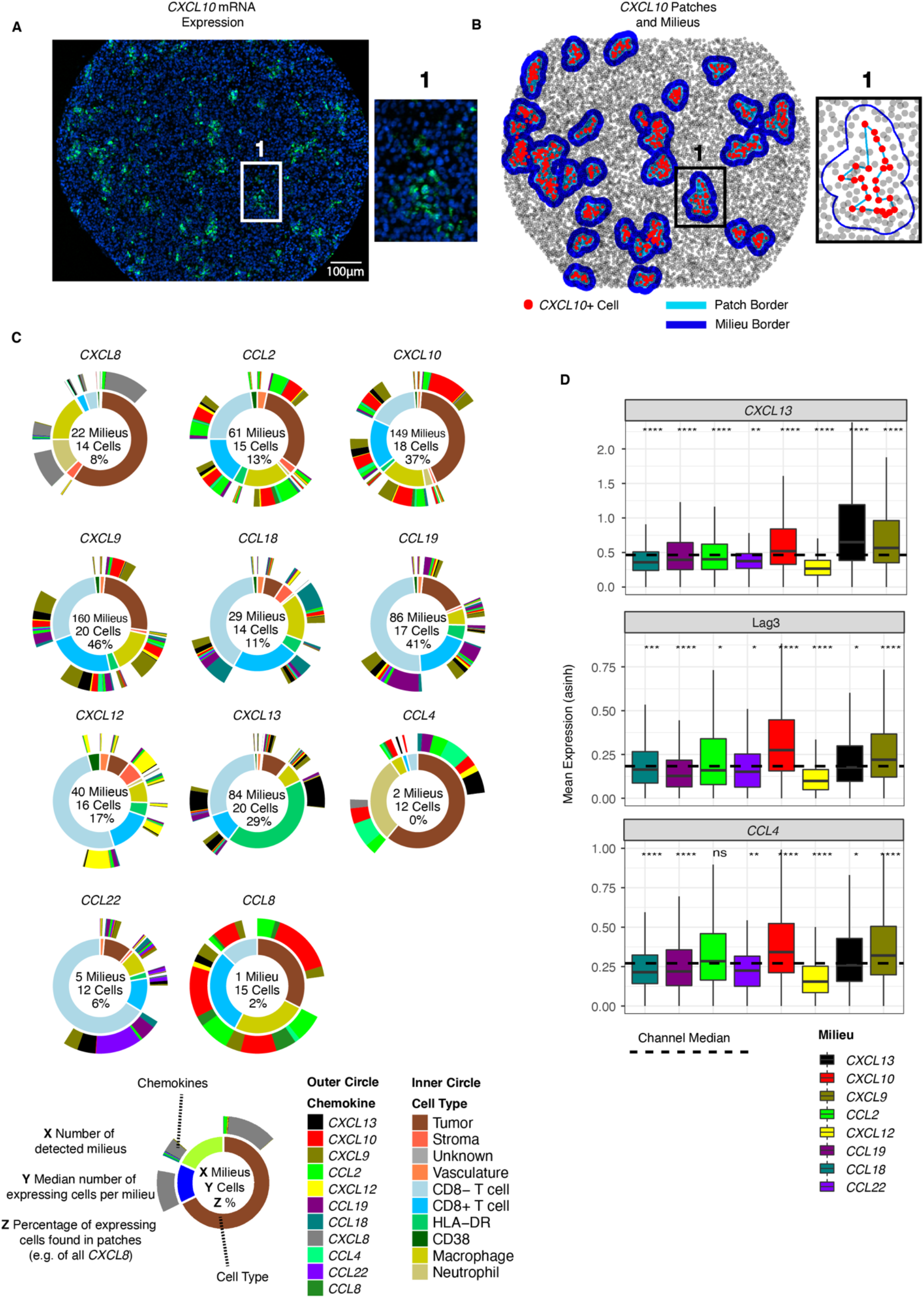
Chemokine-expressing cells form specialized milieus. **(A)** Representative image from a tissue core illustrating that *CXCL10* (green) is often expressed in patches (nuclei are shown in blue). Scale bar, 100 µm. **(B)** Results from the patch and milieu detection algorithm run on the image shown in panel A with one milieu magnified on the right side. *CXCL10*-expressing cells are colored in red. A patch includes only cells that express the marker of interest. Patch borders are highlighted in light blue. A milieu contains all cells of a patch and additionally the surrounding cells. Milieu borders are highlighted in dark blue. **(C)** Sunburst plots summarizing all detected chemokine milieus. The inner circle shows proportions of cell types. The outer circle displays chemokines expressed by the cell types indicated on the inner circle. White space in the outer circle indicates cells that are within a milieu but do not express any chemokine. A numerical summary for each milieu type is given in the circle center. **(D)** Box plots of mean expression of Lag-3, *CXCL13*, and *CCL4* in CD8^+^ T cells for the most abundant milieus. Significance of a statistical test (Wilcoxon, group-comparison against base-median, adjusted using the Benjamini-Hochberg method) for each milieu type is indicated with asterisks.

Local, co-ordinated chemokine expression would be expected to be associated with microenvironments within the tumor that could affect immune cell infiltration and/or function. To further investigate such effects, we went on to characterize the phenotypes and frequencies of cells present within the vicinity of chemokine-expressing cells. We developed an algorithm to detect local accumulations of chemokine-expressing cells (patches) and additionally all the cells within patches and surrounding the patches up to 30 µm away (referred to as milieus) (Fig. 3B). The composition of chemokine milieus are likely to reflect functional consequences of local chemokine secretion.

We compared the overall fractions of cell types and chemokine-expressing cells across all eleven chemokine milieus (Fig. 3C). *CXCL9* and *CXCL10* milieus were the most abundant, with close to half of all *CXCL9*-producing cells and more than a third of all *CXCL10*-producing cells present in milieus. The *CXCL9* and *CXCL10* milieus included cells expressing other inflammatory chemokines at high frequencies and showed the highest frequencies of dysfunctional (*CXCL13*^+^) and recruiter (*CCL4*^+^) CD8^+^ T cells among all milieus, indicating that they represent hotspots of ongoing inflammation. *CXCL10* milieus contained close to 50% of tumor cells indicating that these milieus are at the interface of tumor-immune interactions. Markers for T cell dysfunction (e.g., Lag-3, *CXCL13*, *CCL4*) were particularly highly expressed in *CXCL10* and *CXCL9* milieus in comparison to other milieus (Fig. 3D). Less than 10% of cells that expressed *CCL4*, *CCL22*, and *CXCL8* were observed in milieus, supporting the results found with the local enrichment analysis (Fig. S3B). *CXCL8* and *CXCL12* milieus were most strongly dominated by the expression of the chemokine that defined the milieu.

The *CXCL13* milieus had the highest fractions of HLA-DR^+^/CD163^-^/CD68^-^ cells, which are likely B cells (Fig. 3C). Further, as *CXCL13* milieus also contain CD8^-^ T cells that express *CXCL13*, which may be T follicular helper cells, *CXCL13* milieus are likely enriched in B cell follicles. Of note, since many of our samples were from lymph node metastases, we refer to structures that resemble B cell follicles or TLS within tumors as B cell follicles. Chemokine milieus also differed in their broader cell type compositions. For instance, *CXCL8* milieus consisted mostly of tumor cells, whereas *CXCL12* and *CCL19* milieus showed the highest fractions of CD8^-^ T cells. *CCL18*, *CCL19*, *CXCL12*, and *CXCL13* milieus included at least 75% immune cells indicating that they reside mostly in the stroma. In summary, these analyses allowed us to characterize spatial patterns of chemokine expression and to identify functional chemokine milieus in metastatic melanoma samples.

### Tumor antigen presentation is required for CD8^+^ T cell infiltration

To investigate the differences between tumor phenotypes from hot and cold tumors, we compared the expression of all tumor markers on tumor cells across samples with different levels of T cell infiltration (Fig. 4A, Fig. S3C). Images that contained no CD8^+^ T cells were characterized by a strong reduction in markers associated with antigen presentation (B2M, HLA-DR) and mTOR pathway activity (pS6) and of markers which have been found to be up-regulated upon inflammation in melanoma (PD-L1, Ido1, and H3K27me3) (*39, 40*). Interestingly, we found no difference in β-Catenin, which has previously been associated with T cell exclusion (*41*). The strongest effects were observed for B2M (Fig. 4B), which was confirmed by moderate correlations between the mean expression of B2M in tumor cells with the frequency of CD8^+^ T cells (Pearson’s r = 0.63) and with the fraction of all chemokine-expressing cells (Pearson’s r = 0.57) (Fig. 4C). Thus, in this cohort, antigen presentation by tumor cells and mTOR pathway activity are associated with CD8^+^ T cell infiltration. In tumors without CD8^+^ T cell infiltration, the lack of markers typically up-regulated upon inflammation suggests that these tumors go unrecognized by the immune system.

**Fig. 4.**
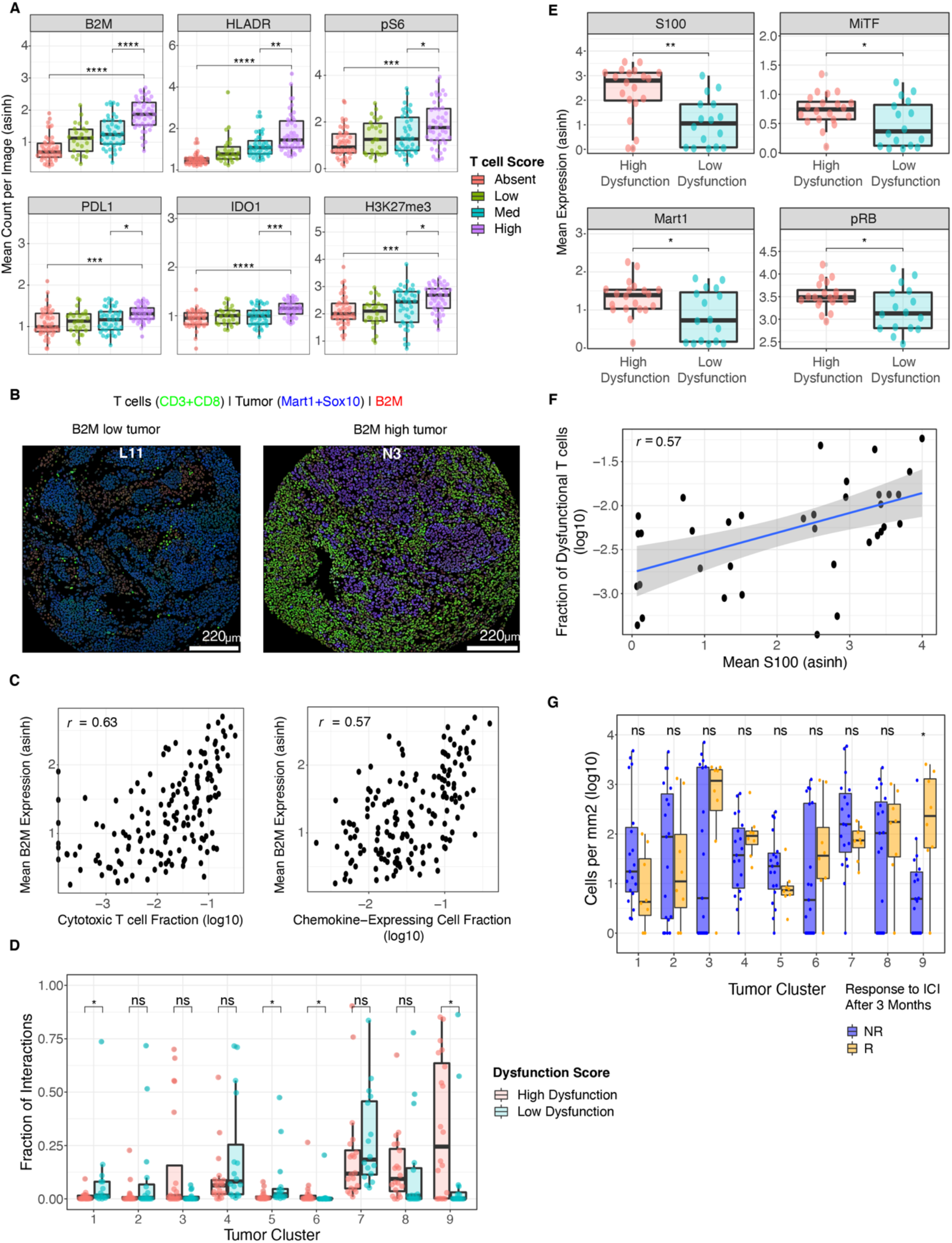
Distinct tumor phenotypes are associated with strong T cell infiltration and T cell dysfunction. **(A)** Box plots comparing the mean marker expression (asinh) in tumor cells of a subset of tumor markers between images grouped by T cell density score. Asterisks indicate significance of statistical comparisons (Wilcoxon, adjusted using the Benjamini-Hochberg method) between images from the “high” density group with the “median” and “absent” groups. One data point indicates one image. **(B)** Cell masks colored by the combined mean expression of CD8 and CD3 (T cells), B2M, and combined Mart1 and Sox10 (Tumor) for two representative images. Color codes are given above the image. Scale bars, 220 µm. **(C)** Left: Scatter plot of the fraction (log10) of CD8^+^ T cells on the x-axis versus mean B2M expression (asinh) on tumor cells on the y-axis. Right: Scatter plot of the fraction (log10) of chemokine-expressing cells on the x-axis versus mean B2M expression (asinh) on tumor cells on the y-axis. One data point indicates one image. **(D)** Box plots comparing the fractions of all interactions of CD8^+^ T cells with different tumor subtypes as defined by clustering between images classified as low- or high-dysfunctional. Significance of statistical comparisons (Wilcoxon, adjusted using the Benjamini-Hochberg method) are indicated with asterisks. One data point indicates one image. **(E)** Box plots comparing the mean expression (asinh) in tumor cells of a subset of tumor markers between images classified as low- or high-dysfunctional. Significance of statistical comparisons (Wilcoxon, adjusted using the Benjamini-Hochberg method) are indicated with asterisks. One data point indicates one image. **(F)** Scatter plot showing the mean S100A1 expression (asinh) on tumor cells per image on the x-axis versus the fraction (log10) of dysfunctional CD8^+^ T cells per image on the y-axis. Linear regression model is shown by blue line; 95% confidence interval is indicated by grey area. **(G)** Box plots comparing the cell density (cells per mm^2^, log10) for tumor subclusters of non-responders vs. responders to ICI (response at 3 months). One data point indicates one image. The p-values were inferred by a statistical comparison (Wilcoxon, adjusted using the Benjamini-Hochberg method) between groups. The responder group included eight images from four patients and the non-responder group included 19 images from seven patients.

### Specific tumor cell phenotypes foster T cell dysfunction

Patients with substantial CD8^+^ T cell infiltration in tumors have better prognosis, but anti-tumor reactivity can lead to T cell dysfunction and exhaustion (*12, 13, 15, 32*). To investigate potential drivers of T cell dysfunction in more detail, we grouped all images with a high CD8^+^ T cell density as either high-dysfunctional or low-dysfunctional based on the fraction of *CXCL13*-expressing CD8^+^ T cells. To avoid inference from adjacent normal lymphoid tissue, images from the tumor margin of samples of lymph node metastases were excluded from these analyses. Images from the high-dysfunctional class were dominated by the presence of *CXCL9*- and *CXCL10*-expressing cells and milieus, whereas images from the low-dysfunctional class showed homogeneous expression of many chemokines (Fig. S4A), further supporting the hypothesis that in tumors with low-dysfunctional CD8^+^ T cells there is ongoing inflammation and anti-tumor responses are active.

We then investigated whether images of the high-dysfunctional class were enriched for interactions between CD8^+^ T cells and specific tumor cell phenotypes. We clustered the protein dataset to obtain fine-grained tumor phenotypes (Fig. S4B): Clusters 4, 5, and 7 were characterized by varying expression levels of Sox10, MITF, and Sox9 and the absence of S100A1. Tumor clusters 3, 6, and 8 were positive for S100A1 and had varying levels of Sox10 and MITF. Tumor cluster 1 was characterized by the sole, strong expression of Sox9, and cluster 2 by the additional expression of MITF and Sox10. Cluster 9 was characterized by strong expression of S100A1 and MITF as well as Sox10 and pS6. We then compared the interaction frequencies of CD8^+^ T cells with cells of each of these tumor clusters between the images of the low- and high-dysfunctional classes. We observed an enrichment of S100A1-expressing clusters 6 and 9 and a depletion of S100A1-negative clusters 1 and 5 in images classified as high-dysfunctional (Fig. 4D). The mean levels of tumor markers S100A1, MITF, Mart1, and pRB were also higher on tumor cells in images classified as high-dysfunctional as compared to those classified as low-dysfunctional (Fig. 4E), although levels of many other tumor markers were similar (Fig. S4C). The strongest difference was observed for S100A1; this was caused by higher frequencies of S100A1-expressing tumor cells in the images classified as high-dysfunctional (Fig. S4D). In agreement with these results, the mean expression of S100A1 on tumor cells per image was moderately correlated (Pearson’s r = 0.57) with the frequency of dysfunctional (*CXCL13*^+^) CD8^+^ T cells per image (Fig. 4F).

The association between dysfunction and S100A1-expressing cell types suggested that these tumor cells might be susceptible to T cell recognition, and that tumors with these cell types are likely to respond to immunotherapy. In support of this, a comparison showed higher proportions of tumor cluster 9 (S100A1^+^) in samples from patients who responded to ICI than in patients who did not respond to therapy (Fig. 4G). In summary, S100A1 expression is increased in tumors with relatively high frequencies of dysfunctional *CXCL13*^+^/CD8^+^ T cells and could potentially serve as a biomarker for response to ICI.

### T cells are required for B cell recruitment

We also carried out a global analysis of cell-cell interactions in images classified as low- or high-dysfunctional. In images classified as low-dysfunctional, almost half of all cell-cell interactions were amongst B cells or BnT cells (Fig. 5A), although B cell frequencies were similar between the two groups of images, and the frequency of regulatory T cells was only slightly (but significantly) higher in images classified as high-dysfunctional (Fig. 5B). The high number of B cell interactions in images classified as low-dysfunctional suggested the presence of B cell accumulations or follicles. Indeed, when we ran the patch detection algorithm on B and BnT cells and then grouped images into four classes (i.e., no B cells, no B cell patches, small B cell patches, B cell follicles), we observed more B cell follicles in images classified as low-dysfunctional (Fig. 5C; p = 0.0498, Fisher-exact test). This shows that the increased number of proximal B cells in images that contained fewer dysfunctional CD8^+^ T cells reflects the presence of B cell follicles rather than more scattered interactions.

**Fig. 5.**
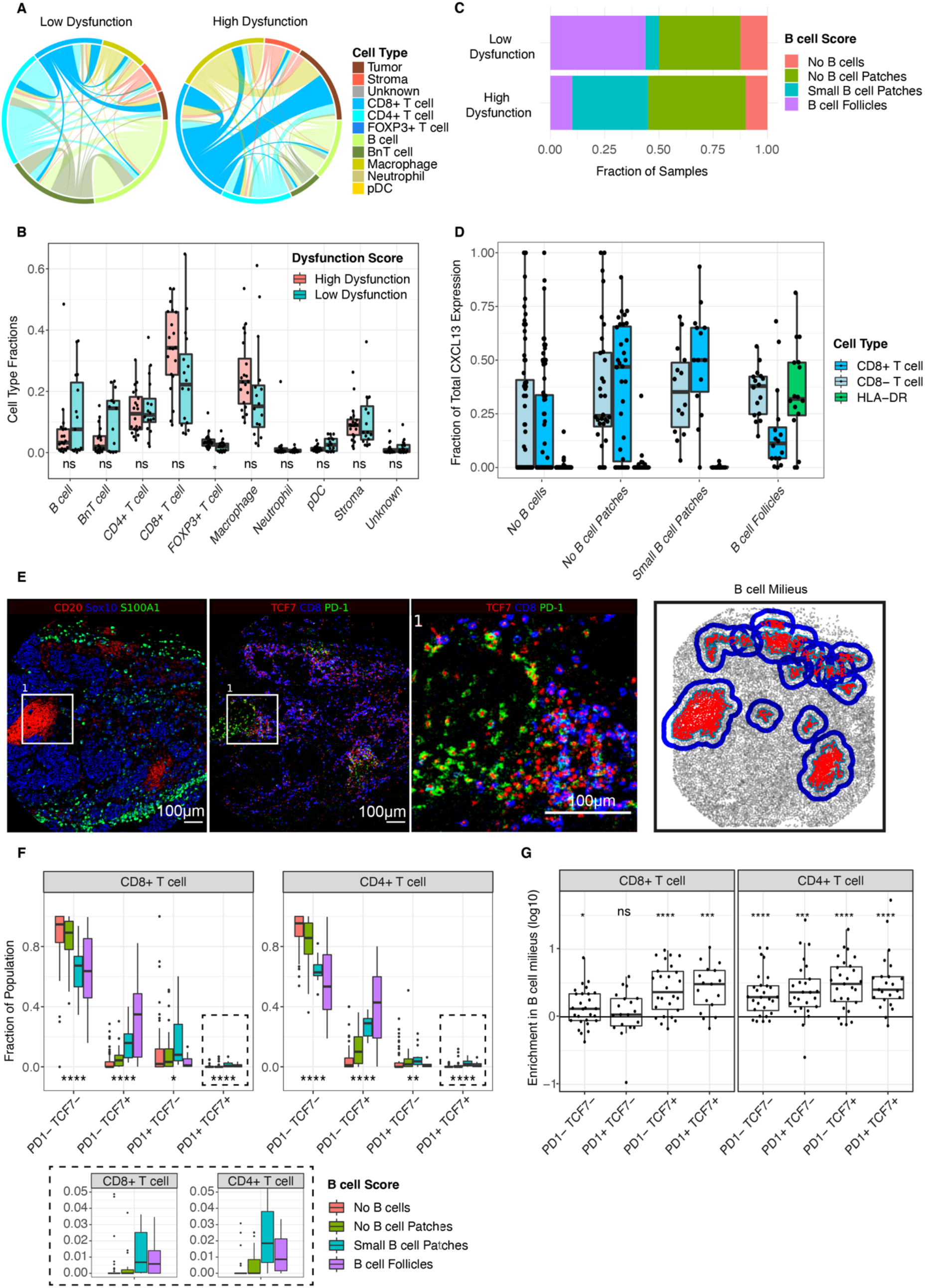
TLS are associated with lower levels of dysfunction and naïve, stem-like T cells. **(A)** Chord diagram summarizing cell-cell interactions for all images classified as low- or high-dysfunctional. Interactions are only shown if they made up at least 5% of all interactions and tumor cell interactions were excluded. **(B)** Box plot comparing cell type fractions for major cell types, excluding tumor cells, between images classified as low- or high-dysfunctional. Significance of statistical comparisons (Wilcoxon, adjusted using the Benjamini-Hochberg method) between groups are indicated with asterisks. One data point indicates one image. **(C)** Stacked bar plots of fractions of samples classified as low- or high-dysfunctional belonging to indicated B cell groups. **(D)** Box plot comparing the fractions of *CXCL13*-expressing CD8^+^ T cells, CD8^-^ T cells, and HLA-DR^+^ cells of all *CXCL13*-expressing cells across the four B cell groups for cells. One data point indicates one image. **(E)** Representative IMC images from staining of one sample with the protein panel colored by different markers. A magnification of the indicated region (white box) from the left and middle images is shown on the right. Marker expression was false colored, and the markers shown are indicated above each plot. A Gaussian blur (sigma = 0.65) was applied. Scale bars, 100 µm. The sketch on the far right shows the results from our B cell patch detection of this image with B cell patches and milieus. B cells and BnT cells are colored in red. Patch boundaries are displayed in light blue and the milieu border is highlighted in dark blue. **(F)** Box plots comparing the fraction of CD8^+^ T cell subtypes (left plot) and the fraction of CD4^+^ T cell subsets (right plot) between the B cell groups. Significance of a statistical comparisons (Kruskal-Wallis, adjusted using the Benjamini-Hochberg method) between B cell groups are indicated with asterisks in every subpopulation. One data point indicates one image. (**G**) Box plots showing the fraction of cell-subtypes inside B cell milieus compared to the rest of the image (log10 scale). This measure was normalized by the relative area of B cell milieus compared to the area of the whole image. Asterisks indicate significance of statistical tests (one sample t-test, µ = 0, adjusted using Holm’s method). One data point indicates one image.

It has been proposed that dysfunctional CXCL13^+^/CD8^+^ T cells are amongst the drivers of B cell follicle formation in lung cancer (*12, 32*) and by CXCL13^+^/CD8^-^ T cells in breast cancer (*42*), potentially through recruitment of B cells. Our dysfunction score is based on *CXCL13* expression in CD8^+^ T cells but does not take into account other *CXCL13*-expressing cell types. We therefore investigated the source of *CXCL13* within tumors corresponding to our four B cell groups in more detail. Our analysis revealed that in images with B cells, but no B cell follicles, the sole sources of *CXCL13* were T cells (Fig. 5D). However, the highest fractions of *CXCL13*^+^/CD8^-^ T cells and *CXCL13*^+^/HLA-DR^+^/CD68^-^/CD163^-^ cells (likely follicular dendritic cells between densely packed HLA-DR^+^ B cells, thus appearing as HLA-DR^+^) were observed in images with B cell follicles (Fig. 5D). We made similar observations for *CXCL13* patches defined in the RNA & protein dataset (Fig. S5A). In images with small *CXCL13* patches, the predominant *CXCL13*-expressing cells are CD8^+^ T cells. With increasing *CXCL13* patch sizes (Fig. S5A, along the DC1 axis) the predominant cells expressing *CXCL13* are CD8^-^ T cells and HLA-DR^+^ cells, confirming our protein-level results. Moderate correlations between *CXCL13* patch size and B cell frequency (Pearson’s r = 0.69) and B cell patch size (Pearson’s r = 0.66) further indicated that large *CXCL13* patches observed in the RNA & protein dataset are likely B cell follicles (Fig. S5B). These data show an association between dysfunctional CD8^+^ T cells and *CXCL13*-expressing CD8^-^ T cells with the presence of B cells and small B cell patches. However, within B cell follicles, *CXCL13* is additionally expressed by what are most likely follicular dendritic cells.

### TLS are enriched in cells with anti-tumor capacity

Naïve-like T cells, defined as TCF7^+^/CD8^+^ cells with or without expression of PD1, have stem-like potential, and their presence is predictive of response to immunotherapy (*12, 15, 16, 32*). TCF7^+^ T cells that did or did not express CD8 often occurred in images with B cell patches or B cell follicles. In particular, PD1^+^/TCF7^+^/CD8^+^ T cells were observed around B cell accumulations (Fig. 5E). We therefore investigated the frequencies of TCF7^+^/PD1^-^ and TCF7^+^/PD1^+^ T cell populations within our four B cell image classes. TCF7^+^ T cell frequencies steadily increased in images ordered from those with no B cells to those with B cell follicles for both CD8^+^ T cells and CD4^+^ T cells with or without PD1 expression (Fig. 5F). The spatial enrichment of TCF7^+^/PD1^+/-^/CD8^+^ over TCF7^-^/PD1^+/-^/CD8^+^ T cells within B cell milieus (Fig. 5G) further supports the importance of TLS with respect to anti-tumor immunity and suggests that these naive-like cells may arise or be primed at TLS sites. In contrast to CD8^+^ T cells, CD4^+^ T cells were generally enriched in B cell milieus (Fig. 5G). We also tested whether the presence of naïve, stem-like T cells was associated with response to immunotherapy in our cohort. Whereas TCF7^+^ T cells appeared to be present at higher levels in responders than non-responders at 3 months, statistical comparisons were not significant (Fig. S5C).

Finally, we performed a correlation analysis between the per-image frequencies of TCF7^+^ T cells and all chemokine-expressing cells to examine potential mechanisms of recruitment of these cells to the tumors. The frequency of *CCL19*-expressing cells was moderately correlated with frequencies of CD4^+^ T cells and TCF7^+^/CD8^+^ T cells but not with TCF7^-^/CD8^+^ T cells (Fig. S5D), suggesting that TCF7^+^ T cells are recruited through CCR7, the CCL19 receptor. TCF7^-^/PD1^+^/CD8^+^ T cell frequencies were highly correlated with frequencies of *CXCL9*-expressing cells (Fig. S5D), suggesting that these cells express CXCR3. A similar picture, albeit with weaker correlations, was observed for *CCL22* and *CXCL12*, suggesting that recruitment of TCF7^+^ T cells may also occur via CCR4 and CXCR4 (Fig. S5D). In summary, we observed naive-like T cells at sites of B cell enrichment within the tumor microenvironment that may arise or be primed at these sites and then induce anti-tumor immune activity.

## DISCUSSION

We have investigated the landscape of chemokine expression in metastatic melanoma using our RNA and protein co-staining approach for multiplex IMC (Schulz et al., 2018). By analysis of the IMC images of samples from 69 patients with tumors from different metastatic sites that ranged in grade from Stage III to IV, we characterized the drivers of immune infiltration and dysfunction in this disease. Why some tumors are immune deserted and others inflamed is a topic of debate, especially since tumors infiltrated by immune cells show higher response rates to immunotherapy (*43*). There was substantial heterogeneity with respect to immune infiltration in the samples from our cohort. We observed that cold tumors are practically devoid of chemokines and showed strongly reduced B2M expression compared to immune hot tumors indicating down-regulated MHC-I presentation, as has been previously reported (*8, 44*). In addition, cold tumors showed lower levels of tumor cell HLA-class-II antigen presentation, of mTOR pathway activity, and of the inflammatory response markers PD-L1 and Ido1, again in agreement with previous reports (*40*). Cold tumors also showed lower levels of the transcription repression mark H3K27me3, which has been shown to be placed by the histone methyltransferase EZH2 to down-regulate antigen presentation upon inflammation (*39*). These data suggest that cold melanoma tumors are dominated by tumor cells that are in “stealth” mode, unrecognizable by the immune system.

Our chemokine expression profiles confirmed known attractive functions of CCL19, CCL22, CXCL9, CXCL10, and CXCL13, which were all analyzed at the mRNA level, toward T cells and B cells (*8, 22–24, 45*). Chemokine expression profiles revealed functional properties of cell types within inflamed tumors, such as *CCL18* expression by macrophages and *CXCL13* and *CCL4* expression by CD8^+^ T cells. We confirmed that *CXCL13*- and *CCL4*-expressing CD8^+^ T cells are strong expressors of Lag-3, suggesting that they have been activated for a sustained period of time and reflect previously described dysfunctional and recruiter cell types (*13, 28, 32*). We also harnessed the imaging capabilities of IMC to study the spatial distribution of chemokines and the cell phenotypes found in chemokine milieus. Similar to findings of a recent study (*26*), *CXCL9* was mostly expressed by myeloid cells, and we often observed CD8^+^ T cells near these *CXCL9*-expressing myeloid cells. Using a novel algorithm to detect accumulations of cells within images, we found that *CXCL9* and *CXCL10* milieus exhibited high expression of markers of T cell dysfunction and exhaustion. In T cell-rich samples, the occurrence of such *CXCL10*- and *CXCL9*-rich milieus with dysfunctional T cells suggests that these are hotspots of inflammation and anti-tumor reactivity.

Multiple states of CD8^+^ T cells exist, from naive-like to cytotoxic to dysfunctional and exhausted, and conversions between these cell states may be possible (*12*). The presence of naïve-like TCF7^+^/CD8^+^ T cell phenotypes is predictive of response to immunotherapy in melanoma and lung cancer (*15–17, 32*). The presence of TLS and B cells are reportedly correlated with response to immunotherapy in melanoma (*8, 18, 46*). Here we discovered that naïve-like, TCF7^+^ T cells are correlated with the presence of TLS in metastatic melanoma. In addition, our IMC analysis revealed that these TCF7^+^ T cells co-occur with accumulations of B cells and show the highest densities around B cell patches and follicles, consistent with recent findings from single-cell analyses (*8*), murine models (*47*), and lung cancer specimens (*32*). Our data thus suggest that the correlations of TLS and TCF7^+^ T cells with ICI response potentially reflect the same underlying events in multiple tumor types.

Our data favor a model in which tumor-reactive, dysfunctional CD8^+^ T cells produce CXCL13 to recruit additional cells of the adaptive immune system to induce a systemic anti-tumor response. We speculate that once a critical mass of B cells is present, the formation of B cell follicles may be initiated, which is further supported and/or sustained by *CXCL13*-expressing follicular dendritic cells. B cell recruitment and follicle formation are likely accompanied by *CCL19*- and *CCL22*-dependent recruitment of naïve and naïve-like T cells that expand upon priming. Furthermore, the fact that we observe these cell types around B cell follicles and associated with *CCL19* and *CCL22* expression suggest that novel clones that have anti-tumor activity may be recruited to the tumor as previously hypothesized (*48*).

Given such a model, it is unclear why certain tumors can, and others cannot, despite the presence of *CXCL13*-expressing T cells, efficiently recruit cells of the adaptive immune system to enter a favorable state for anti-tumor responses. Certainly, tumor phenotypes matter. Interestingly, we found that S100A1 expression is associated with T cell dysfunction and that the presence of specific tumor cells that express S100A1 is predictive of response to immunotherapy, even in our small cohort. Thus, these responsive tumors contained dysfunctional cells, naive-like T cells, and a tumor subtype that may be involved in the response to immunotherapy. Future studies will be necessary to determine whether S100A1, which has so far not been linked to antigen presentation or immunotherapy response, could be used as a biomarker to select patients likely to benefit from ICI.

## MATERIALS AND METHODS

### Experimental model and subject details

#### Biological material

A TMA was prepared from FFPE samples from 69 patients with Stage III and IV metastatic melanoma who were treated at the University Hospital of Zurich under ethics approval KEK-ZH-Nr 2014-0425. For this study, two consecutive (4 μm apart) cuts were processed, stained, and analyzed. The TMA was composed of samples from 14 patients with one biopsy core, 39 patients with two cores, seven patients with three cores, four patients with four cores, and five patients with more than four cores. For those tumors for which we had multiple cores, cores were taken from the tumor border and tumor center.

HeLa cell pellets for the 12-plex validation were obtained from ACD. Overexpression experiments were performed in A431 breast cancer cell lines. Cells were cultured in DMEM (D5671, Sigma-Aldrich), supplemented with 10% fetal bovine serum, 2 mM L-glutamine, 100 U/ml penicillin, and 100 μg/ml streptomycin. A431 cells were only used for overexpression of *PPIB*, and no authentication of the cell line was performed in this study.

### Method details

#### Validation of IMC based RNA detectionfor cytokine evaluation by IMC

To determine the signal intensity range across the twelve channels, channel-specific *PPIB* expression levels were quantified in HeLa cell pellets. To determine the quantitative ability of RNAscope, we compared the median expression level (over three replicates) of 12 housekeeping genes to bulk RNA-seq data from the Human Protein Atlas (*49*). We used normalized expression levels (TMM-normalized, scaled, batch-corrected TPM values) measured in HeLa cells (Expasy Accession CVCL_0030), the cell line evaluated in our experiment.

The influence of subsequent antibody incubation upon the RNAscope protocol was assessed using two replicates of HeLa cell pellets. We quantified the mean expression of 12 housekeeping genes with and without subsequent antibody incubation.

To evaluate crosstalk, a PPIB entry vector from the human ORFeome 8.1 collection (NEXUS Personalized Health Technologies, ETH Zurich) was cloned into a pDEST pcDNA5 FRT TO-eGFP vector (*50*) via Gateway Cloning. A431 cells (ECACC 85090402) were seeded at the density of 50,000 cells per well in a 12-well chambered slide (Ibidi) 24 h before transfection. The cells were transfected with the PPIB expression vector according to the manufacturer’s protocol (Polyplus, jetPRIME^®^ versatile DNA transfection kit). One day after transfection cells were fixed with 10% neutral buffered formalin for 10 min, washed with PBS, and then permeabilized with 0.1% Triton in PBS for 5 min. Following permeabilization, samples were treated with RNAscope Protease III (Advanced Cell Diagnostics) for 10 min at room temperature (1:15 dilution in PBS). Subsequently, cells were washed once in PBS and twice in doubly distilled H_2_O for 2 min each wash. A standard RNAscope protocol (Advanced Cell Diagnostics, RNAscope Fluorescent Multiplex Reagent Kit) was applied starting with single target probe incubation (12 wells = 12 target probes) for 2 h at 40 °C.

The over-expression of *PPIB* was first visually confirmed using a wide-field microscope (Zeiss, Cell Observer) with GFP expression as transfection reporter. Regions with a high density of transfected cells were selected and subsequently ablated using IMC. Although channel spillover effects in IMC are usually minimal, the over-expression of genes enhances the effect to a non-negligible level. Therefore, pixel intensities were spillover corrected using the compensation method described previously (*51*). The degree of nonspecific binding was quantified on the spill-over corrected measurement.

#### Antibody conjugation

The antibodies used in this study are listed in the key resource table. Antibodies (carrier-free) were conjugated to pure isotopes using the MaxPar^®^ labeling kit (Fluidigm) following the manufacturer’s protocol. The antibodies were stored at concentrations up to 500 μg/mL in TRIS-based stabilizing solution (Candor Biosciences) at 4 °C.

#### Staining with RNA & protein panel

Prior to RNA staining, pre-treatment of the FFPE RNA TMA was performed according to the manufacturer protocol (Advanced Cell Diagnostics) for FFPE samples. RNA staining was performed according to the manufacturer protocol (Advanced Cell Diagnostics, RNAscope Fluorescent Multiplex Reagent Kit). Oligonucleotides were conjugated to pure isotopes as described (*29*). Metal-labeled oligonucleotides were then used at a final concentration of 20 nM, diluted from a 10 μM stock with RNAscope HiPlex Probe Diluent (Advanced Cell Diagnostics). After completion of the RNA staining protocol, the sample was briefly washed in TBS. Subsequently, an antibody cocktail was applied and incubated overnight in a humidified chamber at 4 °C. After the overnight incubation, the slide was washed in TBS for 5 min and then stained for 5 min in a 1:100 dilution of 500 μM MaxPar Intercalator-Ir (Fluidigm) in TBS. The slide was then washed for 5 min in TBS, dipped into doubly distilled H_2_O, and subsequently dried using pressurized air flow. The samples were stored until acquisition under dry conditions at room temperature.

#### Staining with protein panel

Slides were first deparaffinized in UltraClear^®^ (3×10 min). After a 10-min wash in UltraClear^®^/100% ethanol (1:1), sections were rehydrated in a graded alcohol series (100%, 90%, 80%, 70%, 50%) for 10 min each, followed by a wash in TBS for 10 min. Antigen retrieval was performed using Tris-EDTA buffer (pH 9.2) for 30 min at 95 °C. After 20 min at room temperature, a blocking buffer (3% bovine serum albumin (Sigma Aldrich), 0.1% Tween-20 (Sigma Aldrich) in TBS) was applied for 1 h at room temperature. The slide was then incubated with antibodies overnight at 4 °C in a humidified chamber. After the overnight incubation, the same steps were applied as described in the RNAscope Staining section.

#### Image acquisition

Images were acquired using a Hyperion Imaging System (Fluidigm). Each TMA core was laser-ablated with a laser frequency of 400 Hz. Cores were randomly selected for acquisition. Five cores could not be acquired as tissue was not visible and most likely lost during sample preparation. A total of 159 images were acquired from each consecutive TMA.

### Quantification and statistical analysis

#### Image processing

Raw data were converted to ome-tiff format and segmented into single cells using the ImcSegmentationPipeline (*52*). TMAs stained with the RNA & protein panel and the protein panel were processed separately as different channels were used to train the pixel classifier described below.

#### Pixel classification

We used Ilastik (version 1.3.2post1) to create pixel probability maps for three classes (nuclei, cytoplasm, background). The class-uncertainty of single cores was extracted to detect cores for which re-training was required. Three cores (H1, I1, I8) were left out in both datasets due to poor image quality. For a subset of cores, pixel classification retraining was performed to reduce uncertainty and to increase the quality of pixel classification.

#### Single-cell segmentation

The generated probability maps were processed to create single-cell masks using the cell image analysis software CellProfiler (version 3.1.9). First, probabilities were histogram equalized (256 bins, kernel size 17), and then a Gaussian filter was applied to enhance contrast and smooth the probabilities. Subsequently, an Otsu two-class thresholding approach was used to segment nuclear masks. Cell masks were derived from an expansion of nuclear masks using a maximum expansion of 3 pixels. Ultimately, CellProfiler was used to overlay single-cell segmentation masks and single-channel tiff images of all measured channels to extract single-cell marker expression means. The mean expression was corrected for channel spill-over using a non-negative least squares method (*51*). Spatial features such as the number of neighboring cells and cell-center coordinates were retrieved. All CellProfiler scripts to recreate the single-cell masks are available at URL (will be released upon publication or request). The output data were then imported into R (version 4.0.3) using the RStudio interface (version 1.3.1093) for further analyses.

#### Cell type identification and definitions

A labeled dataset was generated using the Shiny Application of the R package *cytomapper* (*31*). This application allows labeling of cell populations using multiple gates and inspection of the result on the IMC image to verify the quality of the gating. Gated cells were then downloaded as a *SingleCellExperiment* objects with gating information (min/max information on used markers) stored in the metadata. We defined our cell-type superclasses using the markers available. The markers used for gating and the applied gates for each image and cell type superclass are available at URL (will be released upon publication or request). A random forest classifier was trained on 75% of the labeled data. The model was verified on 25% of the data and subsequently applied to the full dataset. Results from quality control of the random forest classifier are not shown in this publication but are available at URL (will be released upon publication or request).

Since our RNA & protein panel did not contain a probe for *CD4*, we identified CD4^+^ T cells in the RNA dataset as CD3^+^/CD8^-^ cells. CD3^+^/CD8^-^ T cell frequencies in the RNA dataset correlated very highly with CD4^+^ T cells from the protein dataset supporting that CD3^+^/CD8^-^ T cells in the RNA dataset consist of mostly T-helper cells and regulatory T cells (Fig. 1D, dashed box). Similarly, our RNA & protein panel did not directly detect markers of B cells, but HLA-DR^+^/CD68^-^/CD163^-^ cells (referred to here as HLA-DR^+^ cells) correlated very highly with B cell frequencies from the protein dataset suggesting that HLA-DR^+^ cells are highly enriched in B cells (Fig. 1D, dashed box).

#### Detection of chemokine-expressing cells

To reliably identify chemokine-expressing cells, we devised a robust approach that accounts for non-specific probe binding, in which we used the signal obtained with a probe to *DapB*, a bacterial gene not expressed mammalian cells, to infer true positive expression of each chemokine per cell (Fig. S1F). Subtraction of the *DapB* mRNA signal from each chemokine channel for every cell resulted in values that followed a general normal distribution. After scaling these values around a mean of 0 with a standard deviation of 1, p-values for each channel and cell were calculated and adjusted for multiple comparisons using the Benjamini-Hochberg method. A p-value cutoff of 0.01 was applied to all chemokine channels to define cells with significant expression of any chemokine. A binary label was then assigned to each chemokine channel to group the cells into chemokine-producing and non-producing cells. Thus, the expression status of each chemokine in each cell was defined and used as the basis for further analysis.

#### Sub-clustering of cell type superclasses

The classes assigned from the random forest were further sub-clustered into either four or six clusters using only cell-type relevant markers with FlowSOM (*53*) as implemented in the CATALYST R package (*54*).

#### Definition of cell types using manual gating

To manually define cell types, we used a gating approach for the following markers:

● TCF on T cells (CD4^+^ or CD8^+^): mean expression (asinh) > 1.5
● PD1 on T cells (CD4^+^ or CD8^+^): mean expression (asinh) > 1.5
● *CXCL13*^+^ on CD8^+^ T cells: as defined by our chemokine detection method
● S100 on tumor cells: mean expression (asinh) > 3

#### Local enrichment analysis

To identify local enrichments of chemokine expressing cells we used our previously published motif detection version of histocat (*29, 55*). Briefly, motifs were defined by a varying number of cells of the same type (e.g. 4 *CXCL10* expressing) with a maximum distance of 8 µm between cells. P-values were calculated from the comparison of the frequency of motifs per image to the empirical distribution of the number of motifs in the same image after shuffling the cell type labels 1000 times.

#### Patch and milieu detection

In order to detect a local accumulation of a certain type of cell (patch) and their neighbors (milieus), we developed an algorithm in R. The user inputs a *SingleCellExperiment* data container with the IDs for the cells of interest. Additionally required are a maximum distance between cells to be considered as part of a patch and the minimal number of cells by which a patch is defined. The algorithm assigns an ID to cells which fulfill the requirements. The patch border is defined by a concave hull algorithm. To identify the surrounding milieu, a second algorithm expands the patch hull by a user-defined distance value. All cells located within this expanded hull obtain a milieu ID that is identical to the patch ID. For the patch analysis of this dataset, a maximum distance of 25 µm was applied and patch had to consist of minimally 10 chemokine expressing cells. Patches were then further expanded by 30 µm to form milieus. With a median cell diameter of 9.5 µm in our dataset, these expansions corresponded to 2-3 cell diameters.

#### Interaction quantifications

The CellProfiler output from the “measure neighbors” module was used to quantify cellular interactions. For tumor and CD8^+^ T cells, a distance of 8 µm was used to count interactions.

#### Image grouping by T cell score

Images were grouped based on the density (cells per mm^2^ ablated area) of CD8^+^ T cells in the protein dataset. The groups were defined using the following characteristics:

● absent: up to 40 CD8^+^ T cells/mm^2^
● low: 40-100 CD8^+^ T cells/mm^2^
● medium: 100-400 CD8^+^ T cells/mm^2^
● high: >400 CD8^+^ T cells/mm^2^

The scores were derived from the protein dataset and were assigned to the RNA & protein dataset.

#### Image grouping by dysfunction score

Images with high T cell scores were further grouped based on the degree of measured T cell function. Two groups were defined using the following characteristics:

● functional: up to 5% of all CD8^+^ T cells are *CXCL13*^+^
● dysfunctional: more than 5% of all CD8^+^ T cells are *CXCL13*+

The scores were derived from the RNA & protein dataset and were assigned to the protein dataset.

#### Image grouping by B cell score

B cell patches and milieus were defined using our patch detection algorithm. We considered B cells and BnT cells for the detection of patches. The minimal distance between neighboring cells was set to 15 µm. Only patches with at least 20 cells were considered. To define the milieu, the patch border was expanded by 50 µm. Based on the results, we grouped images from the protein data set into four groups:

● no B cells: B cell density of up to 10 B cells/mm^2^ and no patches
● no B cell patches: B cell density > 10 B cells/mm^2^ and no patches
● small patches: at least one patch with up to 250 B cells
● B cell follicles: at least one patch with more than 250 B cells

The scores were derived from the protein dataset were assigned to the RNA & protein dataset.

#### Sample exclusion

To avoid potential biases due to adjacent normal lymphoid tissues, we excluded all images from lymph node margin samples for the calculation of the dysfunction score. Furthermore, lymph node margin samples were excluded for the analysis from Fig. 5D and the B cell follicle analysis shown in Fig. 5F and G. For analyses considering responses to ICI, we only took images from patients with treatment-naive tumors into account.

#### Diffusion maps

Diffusion maps (two diffusion components) were generated on a subset of cells (*CXCL13*^+^) for all markers (asinh transformed) using the scater package (*56*).

#### Statistical testing

Statistical significance was calculated using the pipe-friendly R package rstatix. The statistical test used in each case is given in the figure legend. All p-values were corrected for multiple testing, and the method used is indicated in the figure legend. We used the following convention to indicate significance with asterisks: ns (p>0.1), * (0.1>p>0.01), ** (0.01>p>0.001), *** (0.001>p>0.0001), **** (p<=0.001).

## Supplementary Materials

Fig. S1. Validation of novel 12-plex RNAscope system.

Fig. S2. Chemokine expression and clinical features.

Fig. S3. Chemokine expression across T cell groups and tumor-T cell interactions.

Fig. S4. Protein cell clustering and marker expression comparisons between images grouped by T cell function.

Fig. S5. Chemokine expression and naïve, stem-like T cell associations with B cell milieus.

Table S1. List of reagents.

Table S2. List of software and packages.

## Acknowledgements

We thank the Advanced Cell Diagnostics (BioTechne) team especially Xiao-Jun Ma, Han Lun, and Bingqing Zhang for their continuous support and discussions over the past years. We also thank Natalie deSouza for critically reviewing the manuscript.

## Funding

DS was funded by a Forschungskredit of the University of Zurich, grant no. [K-74419-03-01]. NE was funded by EMBO Long Term Fellowship no. 1194-2019 and the European Union’s Horizon 2020 research and innovation program under Marie Sklodowska-Curie Actions grant agreement No 892225. BB’s research was supported by the European Research Council (ERC) under the European Union’s Horizon 2020 framework, ERC-2019-CoG: 866074 - Precision Motifs and by funding from the Promedica Foundation.

## Author contributions

DS and BB designed and supervised the study. TH and DS performed all experiments and analysis with input from NE. JMG and MPL designed melanoma cohort and prepared the TMA. DS, TH, and BB wrote the manuscript with input from all authors.

## Competing interests

The authors declare no competing financial interests.

## Data and materials availability

Raw data images, mask files for single cell segmentation, and meta data are available under accession number (will be released upon publication or request). The full analysis pipeline was published using the R package workflowr and is available at URL (will be released upon publication or request) (*57*). Separate files are available that enable reproduction of all Figures and Supplementary Figures.

## Notes

### Competing Interest Statement

The authors have declared no competing interest.

