## Supplementary Material for "Multiplexed Imaging Mass Cytometry of Chemokine Milieus in Metastatic Melanoma Characterizes Features of Response to Immunotherapy"

### Affiliations

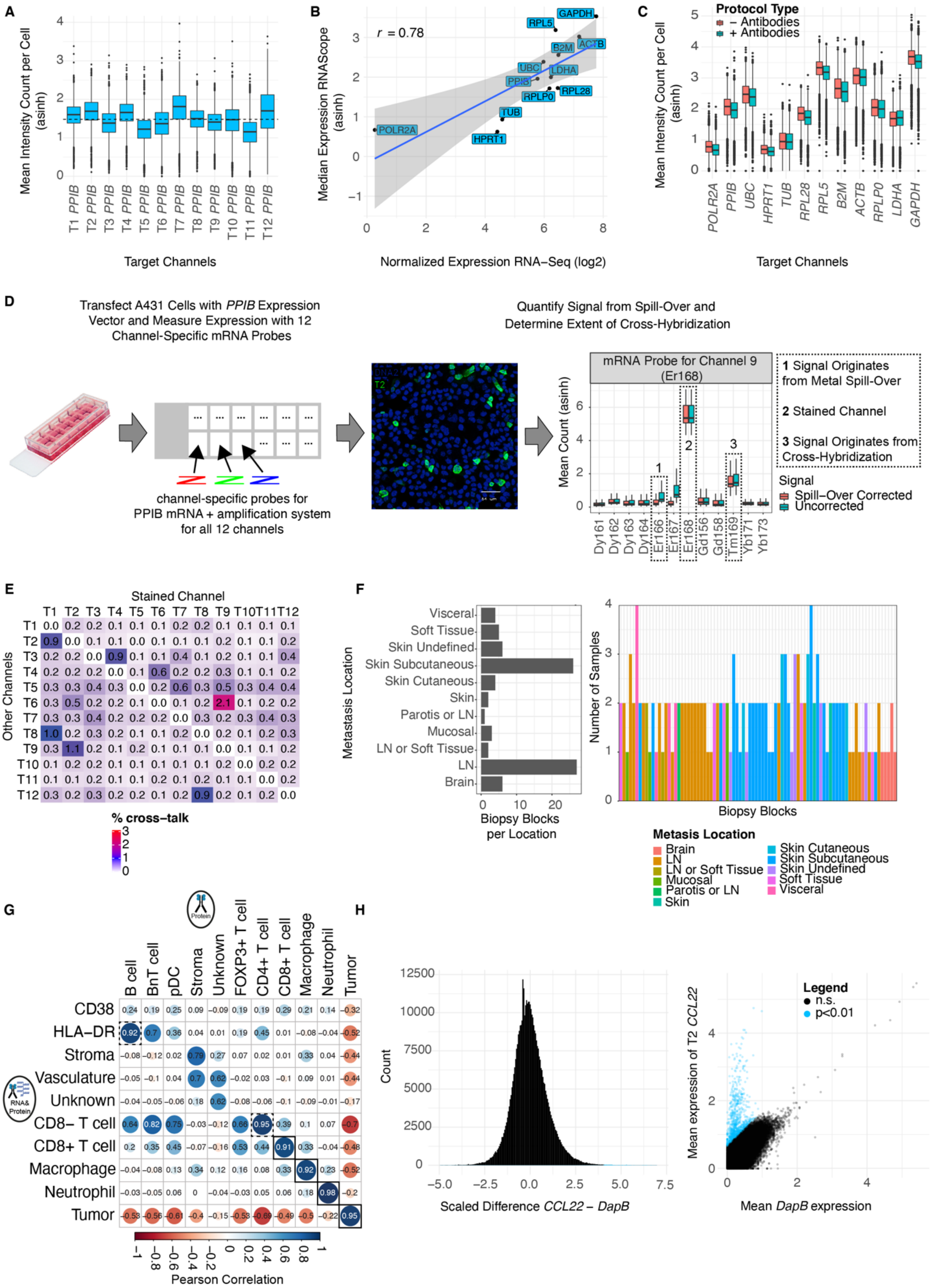

**Fig S1 - Validation of novel 12-plex RNAscope system**

**(A)** Box plot comparing the mean *PPIB* intensity per cell (asinh) measured in 12 different channels for HeLa cells. The dashed line indicates the mean expression across all twelve channels.

**(B)** Scatter plot displaying the normalized RNA-seq expression values (log2) as retrieved from the human protein atlas for 12 housekeeping genes for HeLa cells on the x-axis versus the median expression (asinh) of the 12 mRNAs measured by IMC in HeLa cells on the y-axis (from three replicates).

**(C)** Box plot comparing the mean intensity per HeLa cell (asinh) for 12 housekeeping genes with or without additional antibody staining.

**(D)** Schematic of the experimental workflow to quantify the extent of cross-hybridization events between the 12 RNAscope channels. In each experiment *PPIB* was detected with probes for one channel. During subsequent hybridization steps, amplifiers from all 12 channels as well as metal-labeled probes for all 12 channels were added to detect cross-hybridization between individual channels.

**(E)** Heat map indicating the observed percentage of cross-hybridization amongst the 12 channels.

**(F)** Left: Bar plot of the number of biopsy blocks per location. Right: Bar plot of the number of samples per biopsy block colored according to the metastasis location. LN, lymph node.

**(G)** Pearson correlations of cell type fractions between the RNA & protein dataset and the protein dataset. Correlations highlighted with boxes indicate matched cell types from the two datasets.

**(H)** Representative results from the chemokine-expression detection approach. The bar plot (left) shows the scaled distribution of the real signal (negative control *DapB* signal subtracted from channel-of-interest signal) in all cells. The scatter plot displays the mean negative control signal on the x-axis versus the mean expression for the channel-of-interest on the y-axis. One data point corresponds to one cell (only a subset of cells are plotted for visualization purposes). Significant expression compared to negative control is indicated by light blue coloring in both plots.

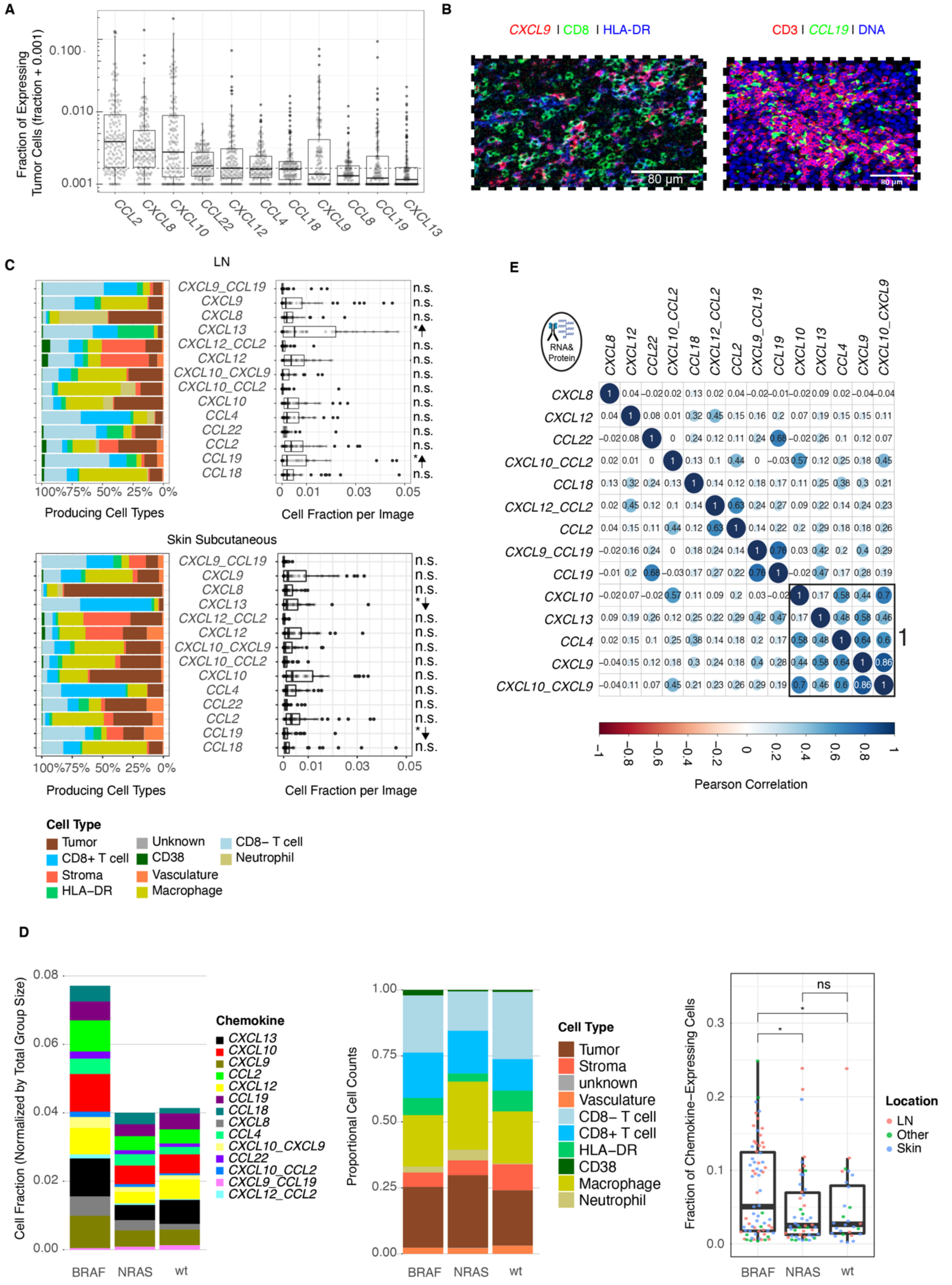

**Fig. S2 – Chemokine expression and clinical features**

**(A)** Box plot comparing the fractions of tumor cells per image expressing indicated chemokines. The dashed line indicates the median fraction of chemokine-expressing tumor cells per image.

**(B)** Left: Representative IMC image of a sample stained with the RNA & protein panel and false colored for *CXCL9* (red), CD8 (green), and HLA-DR (blue). Right: Representative IMC image of a sample stained with the RNA & protein panel and false colored for CD3 (red), *CCL19* (green), and DNA (blue). Scale bars, 80 μm.

**(C)** Left: Stacked bar plots of percentages of cells of indicated types expressing the respective chemokine combination from all images of metastatic lymph node (LN-met) samples (top) and subcutaneous skin samples (bottom). Right: Box plot comparing the fractions of cells per image that express a certain chemokine combination within LN-met samples (top) and subcutaneous skin samples (bottom). Significance of a statistical comparison (Wilcoxon, adjusted using the Benjamini-Hochberg method) between metastasis locations is indicated. Directionality of effect is shown with an arrow.

**(D)** Left side: Stacked bar plot of normalized (by the total number of cells per group) cell fraction for each chemokine combination grouped by mutation. The middle stacked bar plot shows the proportion of chemokine-expressing cell type counts grouped by mutation. The right box plot compares the fraction of chemokine-expressing cells per image for each mutation. Significance of statistical comparisons (t-test, $\mu$ = median fraction over all images, adjusted using the Benjamini-Hochberg method) are indicated with asterisks.

**(E)** Pearson correlations between the frequencies of the top 14 chemokine-expressing cell combinations on the image level.

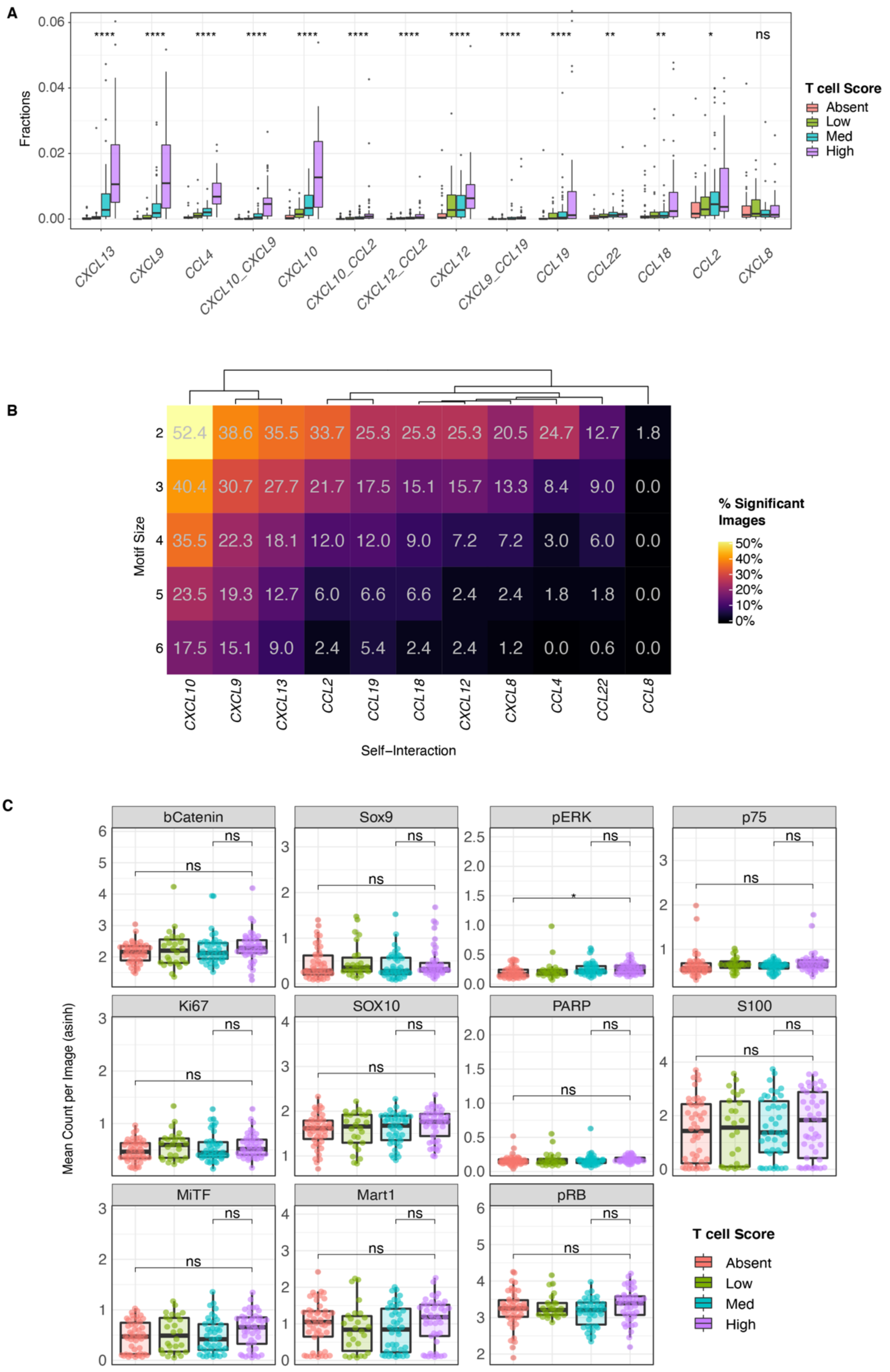

**Fig. S3 – Chemokine expression across T cell groups and tumor-T cell interactions**

**(A)** Box plot comparing the fraction of cells that express a certain chemokine combination per image in the four different T cell density groups. Significance of statistical comparisons (Kruskal-Wallis, adjusted using the Benjamini-Hochberg method) are indicated with asterisks for every subpopulation.

**(B)** Heat map showing the percentage of images with a significant enrichment of chemokine expressing cell motifs of varying sizes (rows) for each chemokine (columns). The percentages are color coded and also given in each tile.

**(C)** Box plots comparing the mean expression (asinh) of a subset of tumor markers in images with indicated T cell density scores. Significance of statistical comparisons (Wilcoxon, adjusted using the Benjamini-Hochberg method) between images from the "high" density group with the "median" and "absent" group are indicated with asterisks. One data point indicates one image.

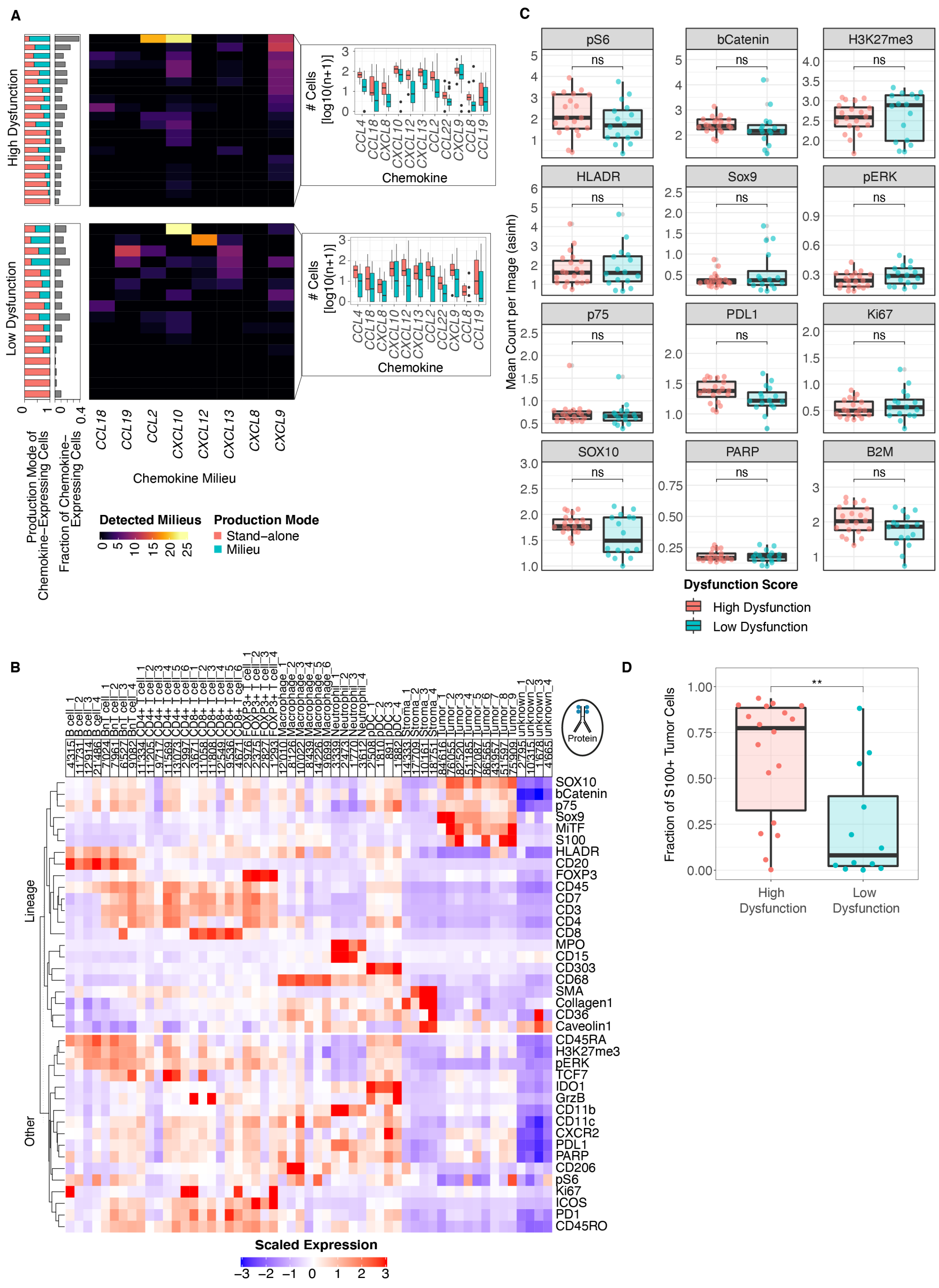

**Fig. S4 – Protein cell clustering and marker expression comparisons between images grouped by T cell function**

**(A)** Clustered heat map split by images classified as low- or high-dysfunctional showing the number of chemokine patches per image. On the left, colored stacked bar plots indicate the fraction of chemokines that are produced in milieus (green) and the grey bar plot indicates the total fraction of chemokine-expressing cells per image. Box plots on the right show the number of cells that express a given chemokine either as part of a milieu (green) or as stand-alone cells (red).

**(B)** Heat map from the protein dataset depicting scaled expression of markers in sub-clustered cell types. Cell counts of each cluster are displayed on top of each column.

**(C)** Box plots comparing the mean expression (asinh) of tumor markers between images classified as low- or high-dysfunctional. Significance of statistical comparisons (Wilcoxon, adjusted using the Benjamini-Hochberg method) between groups are indicated with asterisks. One data point indicates one image.

**(D)** Box plot comparing the fractions of manually gated S100A1$^{+}$ tumor cells (asinh > 3 for S100A1) between images classified as low- or high-dysfunctinoal  Significance of statistical comparisons (Wilcoxon, adjusted using the Benjamini-Hochberg method) are indicated with asterisks.

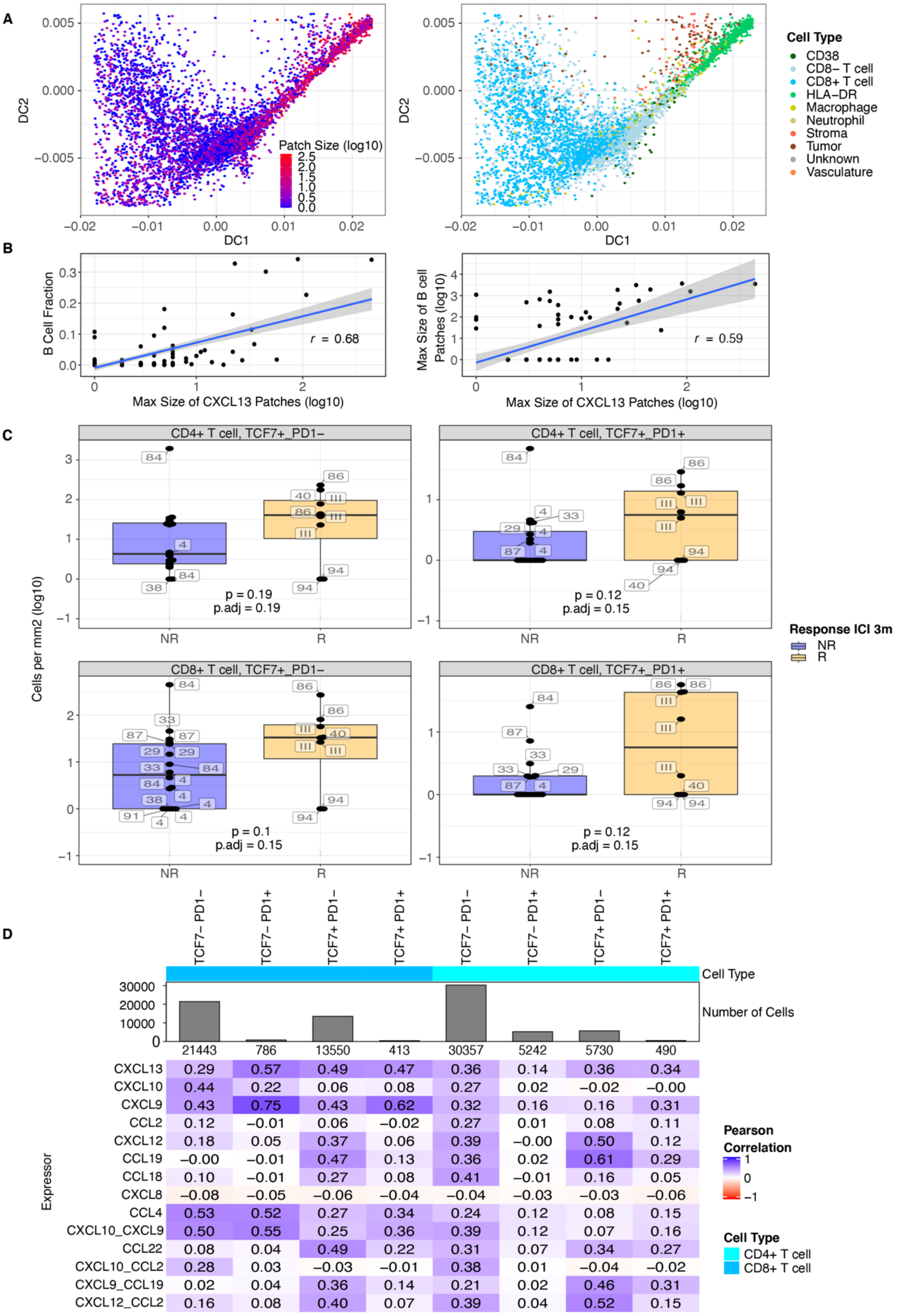

**Fig. S5 – Chemokine expression and naïve, stem-like T cell associations with B cell milieus**

**(A)** Diffusion maps derived from marker expression on all *CXCL13*$^{+}$ cells. One data point indicates one cell. Cells in the left plot are colored by the size of the patch, which ranged from one cell (blue) to several hundred cells (red). Cells in the right plot are colored by their cell type.

**(B)** Scatter plots showing the size (log10) of the largest *CXCL13* patch per image on the x-axis versus the B cell fraction per image (left) or the size (log10) of the largest B cell patch per image (right). The linear regression models are indicated by blue lines; 95% confidence intervals are denoted by grey areas.

**(C)** Box plots comparing the cell densities (cells per mm$^2$, log10) of four T cell subtypes between non-responders (purple) and responders (orange) to ICI (response at 3 months). The responder group included eight images from four patients and the non-responder group included 19 images from seven patients. Labels indicate the patient ID. Significance of statistical comparisons (Wilcoxon, adjusted using the Benjamini-Hochberg method) between groups are indicated with asterisks.

**(D)** Heat map showing the Pearson correlations between the frequencies of cells expressing chemokine combinations and frequencies of TCF and PD1$^{+}$ T expression for CD8$^{+}$ T cells (left four columns) and CD4$^{+}$ T cells (right four columns).

### Table S1 – List of reagents

| **REAGENT or RESOURCE** | **SOURCE** | **IDENTIFIER** |
|---|---|---|
| **Antibodies** | | |
| Vimentin (clone EPR3776) | Abcam | Cat#ab193555;RRID:AB_2814713 |
| SMA (clone 1A4) | Abcam | Cat#ab7817;RRID:AB_262054 |
| CK5 (clone EP1601Y) | Abcam | Cat#ab214586 |
| CD38 (clone EPR4106) | Abcam | Cat#ab176886;RRID:AB_2864383 |
| HLA-DR (clone TAL 1B5) | Abcam | Cat#ab20181;RRID:AB_445401 |
| S100A1 (clone EPR19013) | Abcam | Cat#ab227580 |
| FAP (clone SP325) | Abcam | Cat#ab240989 |
| GLUT1 (clone EPR3915) | Abcam | Cat#ab115730 |
| CD279/PD-1 (clone EPR4877(2)) | Abcam | Cat#ab186928 |
| CD274/PD-L1 (clone 73-10) | Abcam | Cat#ab226766 |
| CD31 (clone EPR3094) | Abcam | Cat#ab207090 |
| Mart1 (clone EPR20380) | Abcam | Cat#ab222483 |
| CD7 (clone EPR4242) | Abcam | Cat#ab230834 |
| CD11b (clone EPR1344) | Abcam | Cat#ab216445;RRID:AB_2864378 |
| MiTF (clone D5) | Abcam | Cat#ab3201;RRID:AB_303601 |

| | | |
|---|---|---|
| CD4 (clone EPR6855) | Abcam | Cat#ab181724;RRID:AB_2864377 |
| Indoleamine-2,3-dioxygenase/IDO (clone SP260) | Abcam | Cat#ab245737 |
| TOX1 (clone NAN448B) | Abcam | Cat#ab256486 |
| Collagen I (polyclonal) | Abcam | Cat#ab34710;RRID:AB_731684 |
| CD11c (clone EP1347Y) | Abcam | Cat#ab216655;RRID:AB_2864379 |
| Myeloperoxidase MPO (polyclonal) | Agilent | Cat#A0398;RRID:AB_2335676 |
| CD3 (clone polyclonal) | Agilent | Cat#A0452;RRID:AB_2335677 |
| p75/CD271 (polyclonal) | Alomone Labs | Cat#ANT-007;RRID:AB_2039968 |
| cleaved PARP (clone F21-852) | BD Biosciences | Cat#552596;RRID:AB_394437 |
| Ki-67 (clone B56) | BD Biosciences | Cat#556003;RRID:AB_396287 |
| Erk1/2 (clone 20A) | BD Biosciences | Cat#612359;RRID:AB_399648 |
| CD134 (clone Ber-ACT35) | BioLegend | Cat#350002;RRID:AB_10639951 |
| CD15 (clone HI98) | BioLegend | Cat#301902;RRID:AB_314194 |
| CCR2 (clone K036C2) | BioLegend | Cat#357202;RRID:AB_2561851 |
| CD45RA (clone HI100) | BioLegend | Cat#304102;RRID:AB_314406 |
| CD45RO (clone UCHL1) | BioLegend | Cat#304202;RRID:AB_314418 |
| Histone H3 (clone D1H2) | Cell Signaling Technology | Cat#4499;RRID:AB_10544537 |
| Beta2-microglobulin/B2M (clone D8P1H) | Cell Signaling Technology | Cat#12851;RRID:AB_2716551 |
| Lag3 (clone D2G4O) | Cell Signaling Technology | Cat#15372;RRID:AB_2798739 |
| pRB (clone D20B12) | Cell Signaling Technology | Cat#8516;RRID:AB_11178658 |
| CD36 (clone D8L9T) | Cell Signaling Technology | Cat#14347;RRID:AB_2798458 |
| TCF1/TCF7 (clone C63D9) | Cell Signaling Technology | Cat#2203;RRID:AB_2199302 |
| CD278/ICOS (clone D1K2T) | Cell Signaling Technology | Cat#89601;RRID:AB_280014 |

| | | |
|---|---|---|
| | | 2 |
| b-Catenin (clone D13A1) | Cell Signaling Technology | Cat#8814;RRID:AB_11127203 |
| S6 (clone D57.2.2E) | Cell Signaling Technology | Cat#4858;RRID:AB_916156 |
| Granzyme B (clone D6E9W) | Cell Signaling Technology | Cat#46890;RRID:AB_2799313 |
| H3K27me3 (clone C36B11) | Cell Signaling Technology | Cat#9733;RRID:AB_2616029 |
| CD279/PD-1 (clone D4W2J) | Cell Signaling Technology | Cat#86163;RRID:AB_2728833 |
| Caveolin-1 (clone D46G3) | Cell Signaling Technology | Cat#3267;RRID:AB_2275453 |
| Sox9 (clone EPR14335-78) | Fluidigm | Cat#3147022D |
| CD163 (clone EDHu-1) | Novus Biologicals | Cat#NB110-40686;RRID:AB_714951 |
| Cadherin11 (clone 283416) | R&D Systems | Cat#MAB1790;RRID:AB_2076970 |
| SOX10 (clone 20B7) | R&D Systems | Cat#MAB2864;RRID:AB_2195180 |
| CD206/MMR (clone 685645) | R&D Systems | Cat#MAB25341;RRID:AB_10890782 |
| CD303 (polyclonal) | R&D Systems | Cat#AF1376;RRID:AB_354762 |
| CXCR2 IL-8 RB (clone 48311) | R&D Systems | Cat#MAB331;RRID:AB_2296102 |
| CD68 (clone KP1) | Thermo Fisher eBioscience | Cat#14-0688-82;RRID:AB_11151139 |
| CD8a (clone C8/144B) | Thermo Fisher eBioscience | Cat#14-0085-82;RRID:AB_11150240 |
| CD20 (clone L26) | Thermo Fisher eBioscience | Cat#14-0202-82;RRID:AB_10734340 |
| FOXP3 (clone 236A/E7) | Thermo Fisher eBioscience | Cat#14-4777-82;RRID:AB_467556 |
| CD19 (clone 6OMP31) | Thermo Fisher eBioscience | Cat#14-0194-82;RRID:AB_2637171 |
| CD45 (clone 2B11) | Thermo Fisher eBioscience | Cat#14-9457-82;RRID:AB_11063696 |
| **Biological Samples** | | |
| Melanoma tissue microarray | University Hospital of Zurich | N/A |
| **Chemicals, Peptides, and Recombinant Proteins** | | |

| | | |
|---|---|---|
| Ir Intercalator | Fluidigm | Cat# 201192B |
| UltraClear | Biosystems AG | Cat# 3905.5000PE |
| Bovine Serum Albumin | Sigma Aldrich | Cat# A3059-500 |
| Tween-20 | Sigma Aldrich | |
| TRIS-based Antibody Stabilizing Solution | Candor Biosciences | Cat# 130 500 |
| **Critical Commercial Assays** | | |
| Spin columns for oligonucleotide purification | Merck Millipore | Amicon Ultra |
| MaxPar® X8 Antibody Labeling Kit | Fluidigm | |
| 12-well chambered slide | Ibidi | Cat# 81201 |
| versatile DNA transfection kit | polyplus transfection | jetPRIME® |
| **Experimental Models: Cell Lines** | | |
| A431 cell line | University of Zurich, Pelkman's Lab | ECACC 85090402 |
| HeLa cell pellets | ACD | Cat# 310045 |
| **Oligonucleotides** | | |
| *CXCL8* channel probe 1 | ACD | custom-made |
| *CCL22* channel probe 2 | ACD | custom-made |
| *CXCL12* channel probe 3 | ACD | custom-made |
| *CXCL10* channel probe 4 | ACD | custom-made |
| *CCL4* channel probe 5 | ACD | custom-made |
| *DapB* channel probe 6 | ACD | custom-made |
| *CCL18* channel probe 7 | ACD | custom-made |
| *CXCL13* channel probe 8 | ACD | custom-made |
| *CXCL9* channel probe 9 | ACD | custom-made |
| *CCL19* channel probe 10 | ACD | custom-made |
| *CCL8* channel probe 11 | ACD | custom-made |
| *CCL2* channel probe 12 | ACD | custom-made |
| *PPIB* channel probes 1-12 | ACD | custom-made |
| positive control mix (12 house-keeping genes) | ACD | custom-made |

| | | |
|---|---|---|
| **Recombinant DNA** | | |
| pDEST pcDNA5 FRT TO-eGFP destination vector | (Couzens et al. 2013) | N/A |
| PPIB entry vector | NEXUS ETH Zurich | Plate: 4530032<br>Row: E11 |

### Table S2 – List of software and packages

| | | |
|---|---|---|
| **Software and Algorithms** | | |
| Ilastik | (Berg et al. 2019) | www.ilastik.org |
| CellProfiler | (Lamprecht, Sabatini, and Carpenter 2007) | www.cellprofiler.org |
| R | R Development Core Team | www.R-project.org |
| RStudio | | www.rstudio.com |
| python | | www.python.org |
| CATALYST | (Chevrier et al. 2018) | www.github.com/HelenaLC/CATALYST |
| workflowR | (Blischak, Carbonetto, and Stephens 2019) | |
| cytomapper | (Eling et al. 2020) | |
| ImcSegmentationPipeline | (Zanotelli and Bodenmiller 2017) | |
| imctools | www.github.com/BodenmillerGroup/imctools | |
| ggplot2 | (Wickham 2016) | |
| SingleCellExperiment | (Amezquita et al. 2020) | |
| ComplexHeatmap | (Gu, Eils, and Schlesner 2016) | |
| reshape2 | (Wickham 2007) | |
| dplyr | (Wickham et al. 2020) | |
| scater | (McCarthy et al. 2017) | |
| dittoSeq | (Bunis et al. 2020) | |
| FlowSOM | (Van Gassen et al. 2015) | |
| sf | www.github.com/r-spatial/sf | |
| concavemen | www.github.com/joelgombin/concaveman | |
| rstatix | www.CRAN.R-project.org/package=rstatix | |
| ggpubr | www.CRAN.R-project.org/package=ggpubr | |

| | |
|---|---|
| ggbeeswarm | www.CRAN.R-project.org/package=ggbeeswarm |

Wickham, Hadley, 2016. *ggplot2: Elegant Graphics for Data Analysis*. Springer.

Wickham, Hadley, Romain François, Lionel Henry, and Kirill Müller. 2020. *Dplyr: A Grammar of Data Manipulation*. https://dplyr.tidyverse.org.

Zanotelli, Vito R. T., and Bernd Bodenmiller. 2017. *ImcSegmentationPipeline: A Pixelclassification Based Multiplexed Image Segmentation Pipeline*. https://doi.org/10.5281/zenodo.3841961.
